# Topographic deep artificial neural networks reproduce the hallmarks of the primate inferior temporal cortex face processing network

**DOI:** 10.1101/2020.07.09.185116

**Authors:** Hyodong Lee, Eshed Margalit, Kamila M. Jozwik, Michael A. Cohen, Nancy Kanwisher, Daniel L. K. Yamins, James J. DiCarlo

## Abstract

A salient characteristic of monkey inferior temporal (IT) cortex is the IT face processing network. Its hallmarks include: “face neurons” that respond more to faces than non-face objects, strong spatial clustering of those neurons in foci at each IT anatomical level (“face patches”), and the preferential interconnection of those foci. While some deep artificial neural networks (ANNs) are good predictors of IT neuronal responses, including face neurons, they do not explain those face network hallmarks. Here we ask if they might be explained with a simple, metabolically motivated addition to current ANN ventral stream models. Specifically, we designed and successfully trained topographic deep ANNs (TDANNs) to solve real-world visual recognition tasks (as in prior work), but, in addition, we also optimized each network to minimize a proxy for neuronal wiring length within its IT layers. We report that after this dual optimization, the model IT layers of TDANNs reproduce the hallmarks of the IT face network: the presence of face neurons, clusters of face neurons that quantitatively match those found in IT face patches, connectivity between those patches, and the emergence of face viewpoint invariance along the network hierarchy. We find that these phenomena emerge for a range of naturalistic experience, but not for highly unnatural training. Taken together, these results show that the IT face processing network could be a consequence of a basic hierarchical anatomy along the ventral stream, selection pressure on the visual system to accomplish general object categorization, and selection pressure to minimize axonal wiring length.

---

Visual object categorization of both non-face objects and face objects inferred from the retinal image is supported by the primate ventral visual stream – a series of hierarchically organized cortical areas (V1, V2, V4, pIT, cIT, aIT) (4–11). We use the phrase “general object recognition” to refer to this set of behavioral abilities. Presently, the best models of the ventral stream neural mechanisms that support this set of behaviors are specific deep artificial neural networks (ANNs) optimized for performance on image categorization tasks (1, 12). While these models can largely explain ventral stream neuronal responses to images, they cannot predict the rich spatial structure of the ventral stream in general and inferior temporal (pIT, cIT, aIT) cortex in particular.

For example, from primary visual cortex to IT cortex, neurons with similar response properties are not scattered randomly within the tissue of each cortical area, but are spatially clustered. In early visual cortex, neurons are primarily organized in the tissue by the location of the visual field corresponding to their retinal-to-LGN inputs (retinotopy), and secondarily by selectivity for low-level stimulus features such as orientation (13), spatial frequency (14), and chromaticity (15). In higher visual cortical tissue, neurons cluster according to selectivity for object categories such as bodies (16), scenes (17), and faces (18).

An important next goal for the field is to build new ANN models that can explain all of these spatial properties, while still explaining the individual neuronal responses. While retinotopy is already partly explained by current deep ANNs (by their initial anatomy), in the present work, we started by asking if new deep ANN models could naturally explain the most salient and well-established spatially-grounded characteristic of high level cortex – the IT face processing network.

The IT face processing network is a constellation of highly robust hallmarks that have emerged from decades of extensive studies in humans and non-human primates on the role of the ventral stream in processing images of faces (18–26). The most detailed neurophysiological studies of face processing have been carried out in non-human primates, and the most salient hallmarks that have emerged from that prior work are: the existence of IT neurons that respond more to face than to non-face objects (“face neurons”) (27, 28), the spatial clustering of face neurons into one to three “face patches” within each level of IT (pIT, cIT, aIT) (23, 29, 30), and the preferential interconnectivity of these face-patches as demonstrated via anatomy and electrical stimulation (31). In addition, it has also been found that invariance to face pose increases along the IT hierarchy (19), paralleling the increase in invariance for non-face objects along the hierarchy (8). Together, these hallmarks are collectively referred to as the IT face processing network (22, 26, 32).

Again, current deep ANN models do not explain all the hallmarks of the face processing network, especially its spatial structure. So what kind of new computational model is needed? The empirically observed spatial organization of neural selectivity in cortical tissue (above) is often considered to be something that might be explained by wiring-length minimization (33). Indeed, wiring-cost minimization inspired the use of self-organizing maps to model the development and geometry of early visual cortex maps for stimulus parameters such as orientation and direction of motion (34, 35). While these models recapitulate the geometry of the topographic maps to an impressive degree, they are limited in their explanatory power. Instead of considering the high-dimensional space of real world images, self-organizing map models assume a drastically reduced space of stimuli generated by a small number of parameters such as retinal position and stimulus orientation. This approach becomes intractable for higher visual areas in which the stimulus parameters of interest are poorly understood. Thus, the kinds of self-organizing maps that have thus far been described do not accurately explain the responses of IT neurons to natural images and they do not explain IT spatial organization.

Here, we sought to overcome these limitations by adopting the underlying theoretical idea (wiring cost minimization), but building upon recent advances in ANN models (1, 12, 36). Notably, that prior deep ANN modeling work has already qualitatively demonstrated the presence of at least some “face neurons” within model IT (36) and more recent studies have demonstrated the existence of face-selective units in deep ANNs (37–39). However, the correspondence of face processing in ANNs and the primate ventral stream has not been tested systematically. Moreover, no prior work has explicitly modeled the spatial organization of IT at the single neuron level. In particular, while a given layer of a deep ANN may have units selective for images of faces (qualitatively similar to monkey “face neurons”), the spatial arrangement of those units cannot be evaluated unless they are embedded in a physical space. In this work, we comprehensively investigate the response properties of face neurons in ANNs using the same measures from the neuroscience literature.

To build new ANN models with topographic structure that can be compared with brain data, we here modify specific deep ANNs by defining the spatial position of each model unit in a two-dimensional space representing the cortical sheet (Fig. 1). We then optimize the synaptic weight parameters of these networks to solve general visual recognition problems (similar to recent prior work) while also minimizing a proxy for wiring cost (old theoretical idea, novel implementation). We refer to this class of models as topographic deep ANNs (TDANNs).

**Fig. 1.**
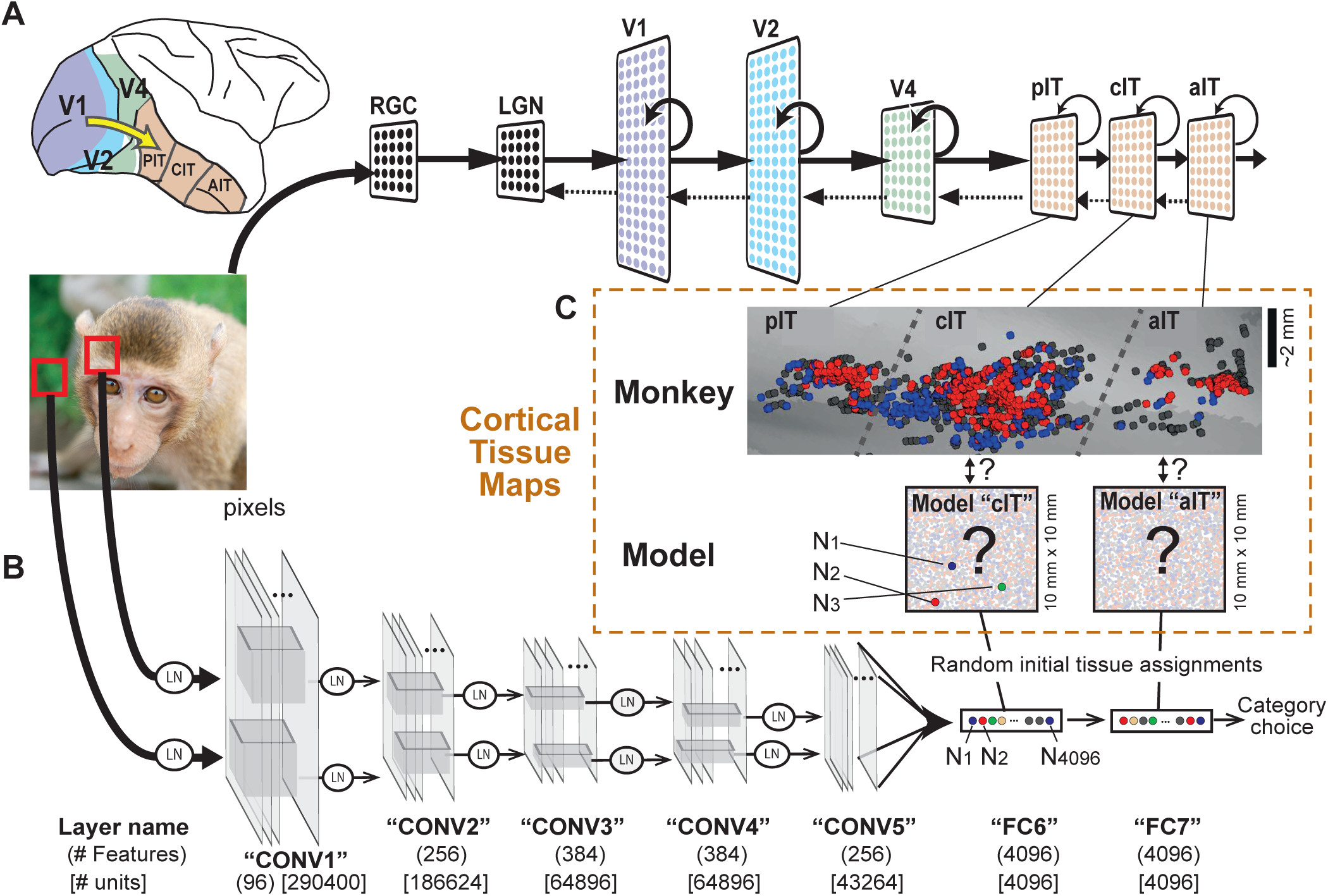
TDANNs as models of the primate ventral visual stream. A) The ventral visual stream is a series of hierarchically organized cortical areas (1). Each stage of visual processing is schematically illustrated on the right as a two dimensional cortical sheet, conceptualizing the spatial arrangement of the output neurons in each area. The dominant feedback, feedforward and recurrent connections are crudely schematized with arrows. pIT, posterior inferior temporal cortex; cIT, central; aIT, anterior; RGC, retinal ganglion cell; LGN, lateral geniculate nucleus (adapted from 1). B) TDANNs are a family of deep artificial neural networks adapted from current deep ANN architectures that are simultaneously optimized to solve a set of visual recognition tasks (“general visual object recognition”) and to also minimize a spatial correlation cost (see Fig. 2). Here, as a base ANN architecture, we use a feedforward only network, AlexNet (2). In the present work, the TDANNs were only required to minimize that spatial correlation cost for neurons within each of its two upper layers FC6 (= model “cIT”) and FC7 (= model “aIT”). It was previously shown that, after general visual recognition training, those two layers of this base architecture (AlexNet) contain artificial neurons that, among all the model layers, are the best predictors of neuronal activity in monkey IT (1). In standard deep ANNs such as this one, the only spatial organization of model units is retinotopy for lower layers (here CONV1 through CONV5), but the spatial layout of the different types of neurons (features) in each of those layers is not specified. Even more problematic, in higher, fully connected layers (here FC6 and FC7), the base model has no spatial organization at all – only a list of model neuron response types (*N*_1_, *N*_2_, etc.) – so it cannot be compared to spatial characteristics of IT. C) To build deep ANNs with predicitons of spatial organization (TDANNs), prior to training, each model unit (*N*_*i*_) in each of the two IT layers (FC6 and FC7) is randomly assigned to a spatial position in an artificial cortical tissue map (a separate 10mm x 10mm map for each layer, see Materials and Methods). During training, the artificial neurons each remain in their initialized position, but their weights (connection strengths with the previous layer) are updated to accomplish correct category assignment of the foreground object in the image (at the output layer, not shown) while also minimizing the spatial correlation costs (see text). Because of this, the TDANN model neurons (*N*_1_, *N*_2_, etc., illustrated in tissue map) are not expected to precisely correspond to the features in a trained version of the base ANN. Gray panel shows data recorded in monkey IT (adapted from 3) with approximate boundaries of pIT, cIT, and aIT. Each dot is a recording location and color indicates relative selectivity for faces (red) vs. non-face object (blue). Note one dominant face patch in each sub-region of IT.

After successfully building specific, functioning TDANNs, we tested whether they recapitulate well-established features of the primate IT face processing network. Indeed, we find that TDANNs naturally reproduce the key empirical hallmarks that together define the face processing network: the existence of face-selective units in model IT (27, 28); the topographic arrangement of those face-selective units that, without parameter tuning, naturally have the same spatial extent as primate face patches (40, 41); stronger fMRI-detectable selectivity of face patches than for most other object categories (25, 42); connectivity between face-selective regions at different stages of the visual hierarchy (31); and the emergence of viewpoint invariance throughout the IT face processing network (19). Further, we empirically find that these phenomena emerge when the networks are optimized for all tested types of naturalistic experience, but not for highly unnatural experience. Overall, these results show that the IT face processing network could be a consequence of a basic hierarchical anatomy along the ventral stream, selection pressure on the visual system to accomplish general object recognition, and selection pressure to minimize neural wiring length.

## Results

### TDANN models retain high performance on general visual recognition while also reducing wiring costs

To test our overarching hypothesis (that the IT face sub-network might be an emergent consequence of the visual system to solve general visual recognition while also minimizing wiring costs), we first had to engineer deep ANN models that could successfully implement these underlying goals. Specifically, we asked if we could build deep ANN models that solve two challenges: general visual recognition and wiring cost minimization. Given the computational difficulty of directly minimizing wiring costs, we implemented a spatial correlation rule as a proxy of that cost within specific layers of the ANN model (Fig. 2; see Materials and Methods for details). In building these models, our underlying rationale was that in any hierarchical anatomy (i.e., pIT, cIT, aIT), there is evolutionary/developmental pressure to minimize general visual recognition errors and minimize wiring cost, and that wiring cost minimization pressure can be approximated as a pressure to have neurons with similar response profiles located physically nearby each other, which is implemented as a spatial correlation rule. The one free parameter of the spatial correlation rule (i.e., the fall off of response similarity per mm of cortical tissue) was derived from actual monkey data (Fig. 2A). Importantly, this scaling parameter was not derived from IT face patches. Instead, it is known to be a general phenomenon of IT cortex (43, 44; we re-confirmed this in Fig. 2A with 2B where all face neurons are excluded from the analysis).

**Fig. 2.**
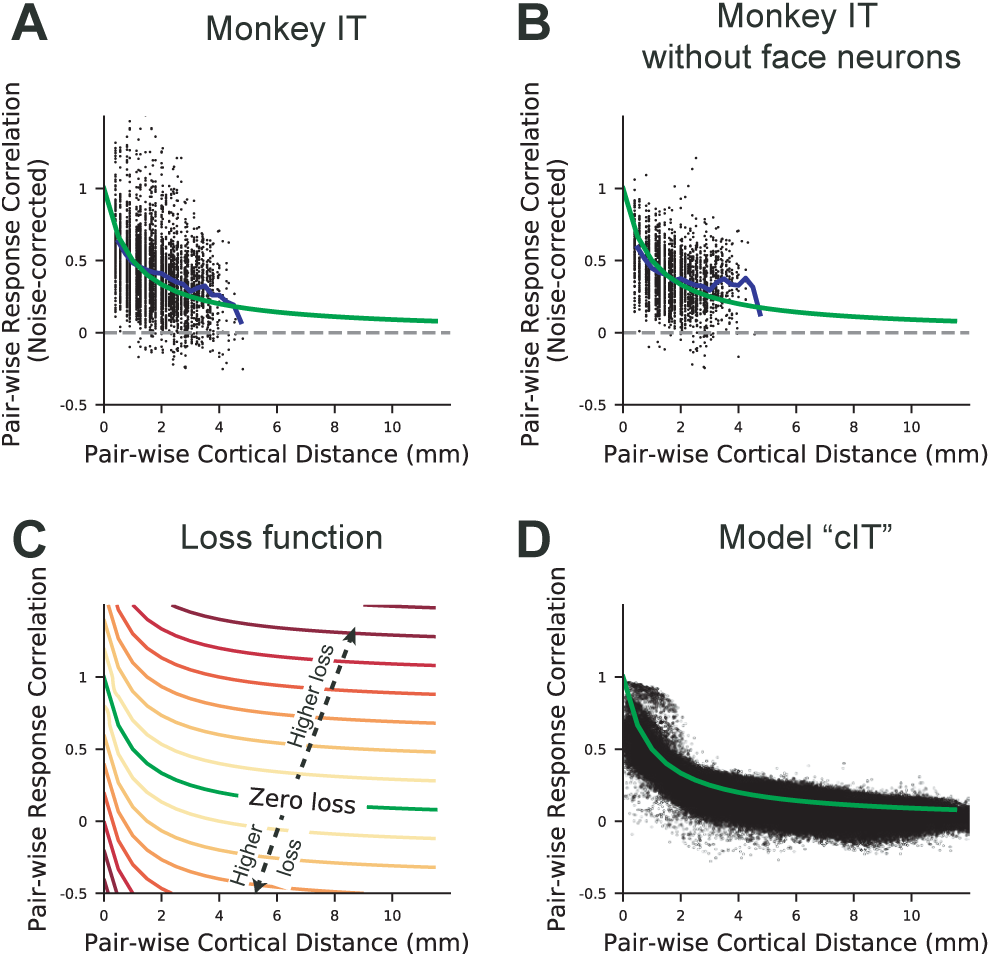
Empirically observed spatial correlation of IT neuronal response profiles and spatial loss function for TDANN models. A) The response profile of each IT neuron is a list of measurements each corresponding to the mean response of one of 5760 images (mean over ∼50 repetition tests of that same image; 47). Here, each black dot is the Pearson correlation of the response profiles of a pair of neuronal sites recorded in monkey IT, plotted against the physical distance between those two sites (measured approximately tangentially to the cortical surface). To remove the effects of noise and repetition variability, the correlation values were normalized by the split-half reliability of the neurons across image repetitions of the images. The blue line is the mean over each 0.25 mm wide spatial window. B) Analysis of the same IT data, except that all face-selective neurons (*d*′ > 0.65) were removed, demonstrating that the dominant spatial correlation profile from (A) is a general phenomena of IT cortex and is not simply a result of face patches. In all panels, the target spatial correlation profile for the TDANN models is overlaid as a green line (referred to as the “zero loss” line in C). C) The target response correlation spatial profile for the model units in model cIT and in model aIT. The zero loss curve was parameterized to approximate the actual IT response correlation spatial profile (A, see Materials and Methods for details). A TDANN model incurs the lowest overall spatial loss when all its pairs of units fall exactly along the green curve – pairs of model units that fall off that profile in either direction incur higher loss (colored lines). D) Data from one examplar TDANN model cIT after training to optimize for visual recognition and for this spatial loss. Each black dot represents a pair of model cIT units.

During the training of each TDANN model, we enforced the spatial correlation rule for units in the model’s final two fully-connected (FC) layers (layers FC6 and FC7). For the base architecture used here (AlexNet, 2), these two layers were already known to be the best predictors of neuronal activity in IT after image general visual recognition training (1). Because most of the relevant monkey data is from cIT and layer FC6 is a better predictor of cIT than FC7 (Fig. S10), we focus on layer FC6 (model cIT), but we include results from layer FC7 (model aIT) where comparison between adjacent layers is warranted (e.g. Fig. 5 and S8). Hereafter, we refer to layer FC6 as model “cIT” and FC7 as model “aIT”. For the baseline TDANN training, we optimized the network to solve an interleaved set of 914 visual image categorization tasks that included just one face category (our operational definition of “general visual recognition”). These categories were constructed using a dataset consisting of naturalistic images, subsampled from the non-vehicle-related ImageNet (45) images and the Labeled Faces in the Wild (LFW) images (46). We refer to this as the “crudely ecological” dataset. We trained twenty such TDANN models, each of which has the same base architecture but randomly, and thus differently, initialized parameters.

**Fig. 3.**
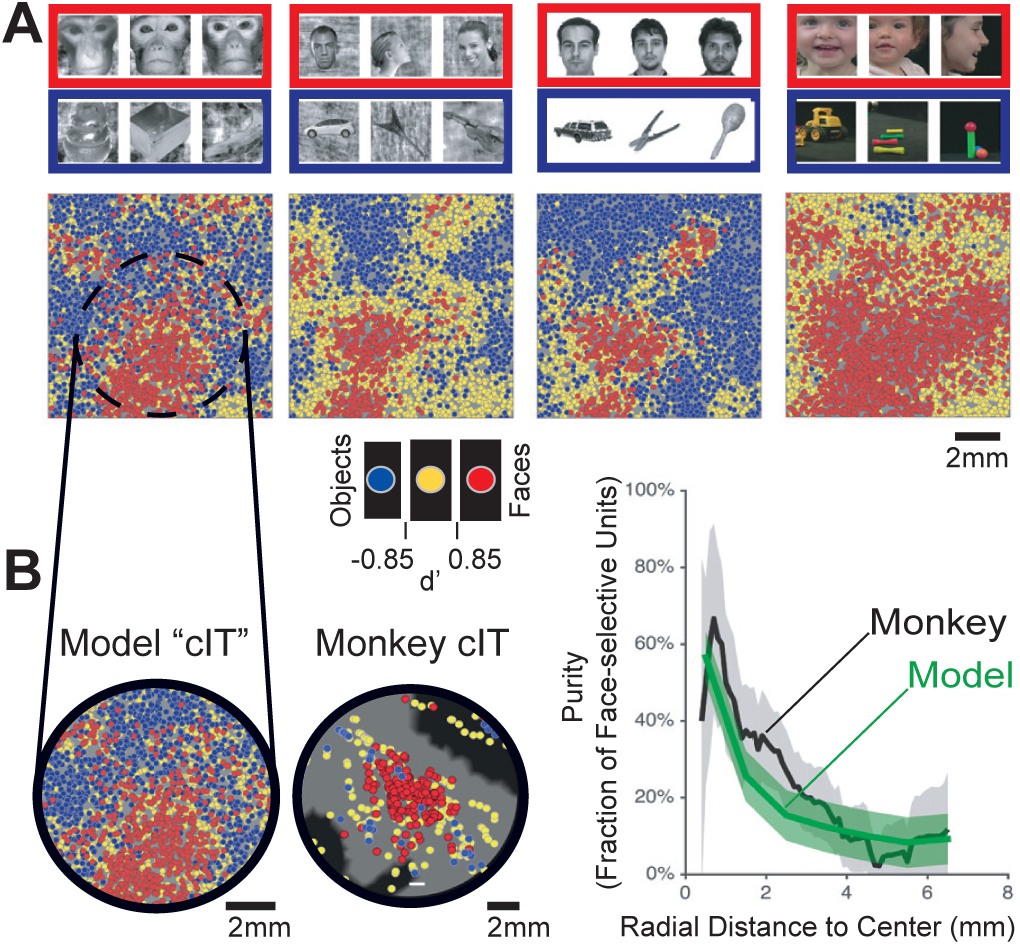
Face selectivity maps from macaque cIT and trained TDANN model cIT. A) Spatial distribution of face-selective responses in the TDANN cIT layer of a single representative TDANN model, as assessed by four different face selectivity test image sets (40, 25, 42, 41, from left to right). In each case, face-selective units (*d*′ > 0.85) are shown in red; non-face object-selective units (*d*′ < −0.85) are shown in blue; units that are not strongly biased to either of those categories (−0.85 *≤ d*′*≤* 0.85) are shown in yellow. When different test sets are used to probe the responses of the model neurons, its face selectivity map is not identical, yet the locations of the dominant clusters of face-selective units remain largely the same. B) An enlarged view of the dominant model cIT face-patch in panel A (left) and actual neurophysiology data from the monkey middle face patch (cIT; middle, 40) shown at identical spatial scales. Rightmost panel shows purity of face selectivity as a function of distance from the center of the dominant cIT face patch in monkey (black, 40) and TDANN models (green; here an average of 20 TDANN models with different weight initializations prior to training; purity curves of individual models can be found in Fig. S4). The gray shaded region for macaques and green shaded region for the models indicate the standard error of mean (SEM). All scale bars, 2mm.

**Fig. 4.**
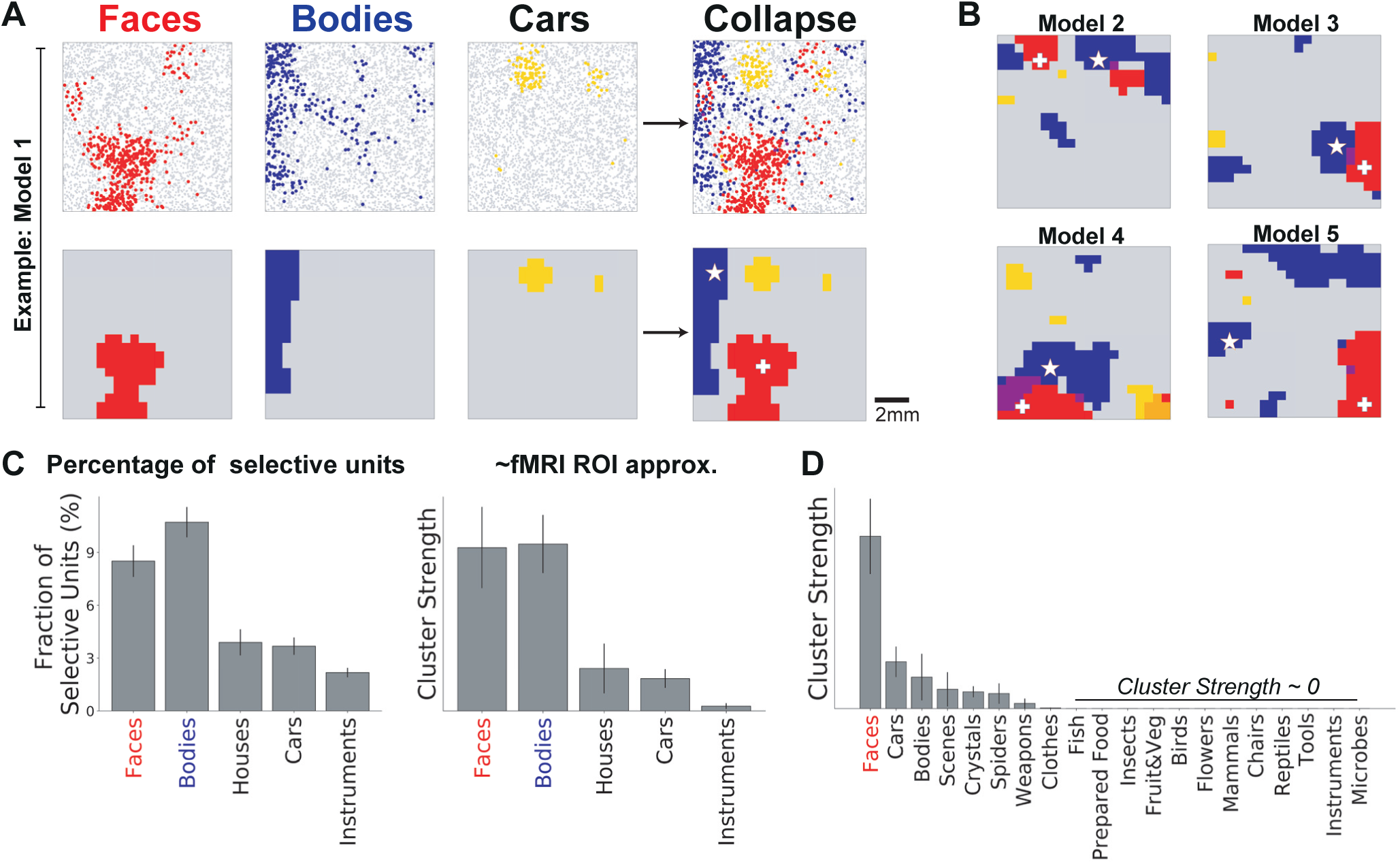
Category selectivity of TDANN model cIT. A) Model unit category selectivity maps (upper; each dot is a model unit) and simulated fMRI maps (lower) in a representative TDANN model. All panels are the same 10mm x 10mm cIT tissue map from a single TDANN model. The left three maps show selectivity measured in response to test images of faces (red), bodies (blue), and cars (yellow), respectively (see Materials and Methods). The rightmost plots show the superposition (collapse) of the three preceding plots. Units that were selective for both faces and bodies (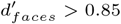 and 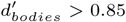) are shown in purple. Each simulated fMRI map is an estimate of the fMRI map that would result from the same image test sets for faces, bodies, and cars, given the spatial arrangement and selectivity patterns of the neural units in the model (see Methods for details). The white-colored plus and star markers indicate the locations of peak selectivity for faces and bodies, respectively. B) Additional simulated fMRI collapse maps from four other TDANN models, each with a different random initialization. C) Left: Mean percentage of category-selective units (*d*′ > 0.85) over all TDANN model cITs (n=20). Right: mean cluster strength of category-selectivity over all TDANN model cITs (n=20). This measure approximates standard fMRI ROI analysis by first selecting the largest cluster in each layer from a subset of test images, and then testing with new images (see Materials and Methods). D) Same as C, but using a different standard fMRI image test set with more categories (16). Categories under the bar yielded barely measurable clusters. Except in (D), all analyses were done with the test images from (25). In C and D, error bars indicate the SEM over the 20 TDANN model runs.

**Fig. 5.**
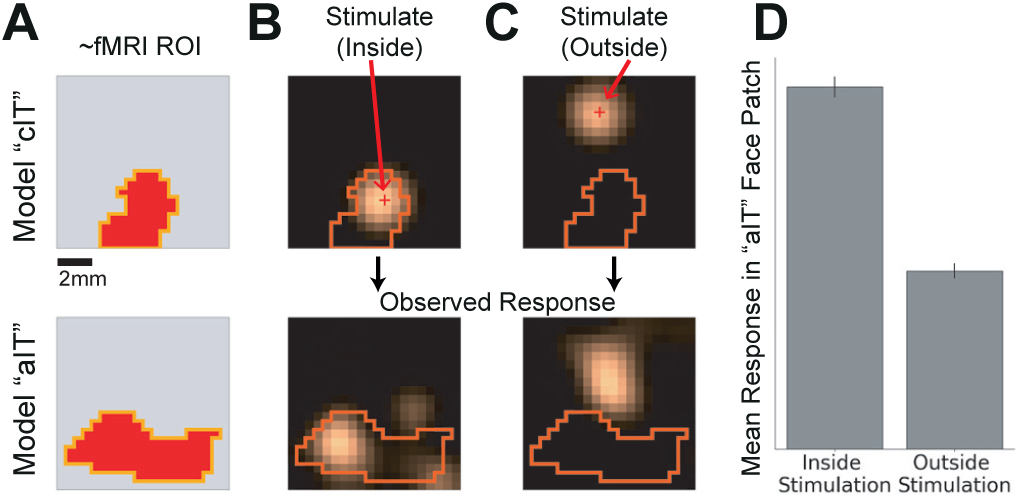
Simulated electrical stimulation inside model cIT face patches leads to fMRI- detectable activation of model aIT face patches. A) Maps of face-selectivity in model cIT (upper) and aIT (lower) of an example TDANN model (located as described in Materials and Methods). Orange lines demarcate the boundaries of face patches and are replotted in B and C. B) Electrical stimulation inside of the model cIT face patch (upper) and the observed responses in model aIT (lower). See Materials and Methods for details on simulated stimulation. The red plus marker in the model cIT map indicates the simulated stimulation site and the color map represents the stimulated activation of units around the stimulation site in cIT (upper) and in aIT (lower). C) Stimulation outside of the face patch in model cIT does not lead to an appreciable increase in the activity of model aIT face patch units. D) Quantification: Mean activation of the model aIT face patch units for stimulation inside (left) and outside (right) of model cIT face patch (mean across 20 randomly-initialized TDANN models; in each case, all face-selective regions in cIT and aIT were identified and its boundary was defined as described in Materials and Methods). Error bars indicate the SEM.

We found that, compared to the non-topographic control ANN models (which had the same architecture without a spatial cost), each TDANN model achieved nearly the same level of general visual recognition performance on the held out validation images (Fig. S1D) while each model also adhered to the spatial correlation rule (Fig. 2). The degree to which the TDANNs adhered to this rule can be visualized in Fig. 2D, which shows the model cIT units from one exemplar TDANN model. In this case, the closer the model units are in the “cortical tissue”, the more similar their response patterns were as encouraged by the spatial correlation rule. Interestingly, the model cIT units did not perfectly align with this rule and showed a similar degree of divergence from the rule (black dots spread around green line in Fig. 2D) as the actual IT neurons (Fig. 2A,B) even though the degree of variability was not directly dictated by the rule.

We also asked, do TDANN models actually have lower wiring costs? Indeed, we found that, under some simple assumptions about total wiring length needed to implement fully trained models, the TDANN models do indeed have lower wiring costs than regular fully trained ANN models of the same base architecture (see Supplementary Information for details). Because this is just one of many ways to analyze wiring costs and it is challenging to optimize for such costs directly, this line of work will be continued in follow-up studies (e.g. Jozwik et al., unpublished). In the present work, our main result is that the addition of a spatial correlation rule within each level of the IT hierarchy is sufficient to reproduce the hallmarks of the IT face network (below) and it also tends to reduce overall network axonal wiring costs.

### TDANN category-selectivity maps mirror the spatial organization of primate IT cortex

In human and macaque higher visual cortex, “face neurons” and “body neurons” that respond more strongly to images of faces or bodies than to non-face and non-body objects are not scattered randomly across IT cortex, but are instead spatially organized. The presence of face units in current (non-topographic) deep ANNs is a topic of active research, but several studies suggest that current baseline ANN models do indeed contain face neurons (37–39), and we confirm that here with methods that operationally match those used in primate and human neuroscience experiments (Fig. S2). However, those current ANN models cannot explain the spatial organization of face-selective units because they are non-topographic. Here, we ask if the class of TDANN models predicts not only the presence of face units, but their spatial organization as well.

We first investigated the spatial pattern of category selectivity over TDANN model IT neurons by evaluating each model’s neuronal responses to images of faces, bodies, and other objects from four standard image test sets (25, 40–42) that have previously been used to localize category-selective regions in humans and macaques (Fig. 3, 4). We investigated responses to these images in the TDANN model cIT (see Materials and Methods). As detailed below, evaluation of the resultant category preference maps suggests close correspondence to several of the key spatial organizational features reported in humans and macaques when measured with similar or identical image sets.

#### TDANN units selective for faces and bodies cluster into patches, while other standard categories do not form strongly selective clusters

We first investigated the spatial pattern of face selectivity, where a unit was defined as face-selective if the *d*′ of its response to images of faces vs. images of non-face objects was greater than a threshold of *d*′ > 0.85. This threshold was selected to correspond to the threshold of *d*′ > 0.65 used in (40) after simulating the addition of Poisson noise to model responses (see Materials and Methods for details). We found that face-selective units in the TDANN model cIT were spatially clustered in a similar fashion to the clustering of face-selective neurons in the macaque middle face patch (23, 40). Fig. 3 shows this for one example TDANN model (using one initial seed for training) and Fig. S3 shows more from differently initialized TDANN models. We also found that when different operational definitions of face selectivity were used (i.e., different image sets comprised of monkeys, human children, and human adults), the dominant cluster of face-selective neurons was localized to the approximately the same region of the map (Fig. 3A). The reliability of the spatial pattern of face selectivity over all differently initialized TDANN models and across different stimuli used to measure face selectivity is provided in Fig. S14A (see Supplementary Information for details).

It should be stressed that even though TDANN cIT and aIT layers were not explicitly constructed to contain face patches, they nearly all exhibited face patches (Fig. 3, S3; quantified below). Nevertheless, because these model layers were constructed to produce neural response selectivity maps that tend to be spatially smooth (see Fig. 2 and Materials and Methods), perhaps it logically follows that TDANN cIT and TDANN aIT *must* contain face patches? It does not, as evidenced by the brain: some, but not all object categories, show fMRI-level spatial clustering. In particular, besides the category of faces, fMRI studies have thus far found reproducible regions of interest for categories of bodies, scenes, color, and words (16, 17, 25). Moreover, in some cases, such clustering has been verified at the single neuron level in monkeys (43, 48, 49). Importantly however, fMRI studies also show that other standard object categories (e.g., cars, musical instruments, rocks, etc.) do *not* yield observable clusters in humans (42, 50, 51). Thus, we asked, do TDANN cIT and TDANN aIT predict these same trends? To answer this question, we performed contrast analyses that mimicked fMRI experiments on each model using a functional localization stimulus set including images of faces, bodies, houses, cars, and musical instruments (25).

Fig. 4A shows maps of selectivity (*d*′ > 0.85) for three of the categories in the functional localization stimulus set: faces (red), bodies (blue), and cars (yellow), for one example TDANN model cIT. Maps of selectivity to the other categories and for other TDANN model cITs are included in the Supplementary Information (Fig. S6) but omitted here for clarity. To more directly compare the topology of category selectivity in our models to results obtained with fMRI, we simulated high-resolution (0.5mm isotropic) fMRI maps by smoothing each model’s spatial pattern of neural activation for each image with a 1mm Gaussian kernel (Fig. 4A, lower) and then recomputed category selectivity on a per-voxel level, where selectivity was defined by a threshold: *t* > 10. Qualitatively, we observed that there were more face- and body-selective units than car- and instrument-selective units in both the native neural maps (Fig. 4A upper), and the fMRI-simulation voxel maps (Fig. 4A lower, 4B and S6). This result was robust across maps from models with different randomly-initialized weights: we observed clear face- and body-selective patches in 18 of the 20 randomly-initialized models, where a patch was identified as the presence of at least 10 spatially connected voxels selective for the category.

Moreover, quantitative analyses support our qualitative observations. We found more units selective for faces and bodies than to any other category (Fig. 4C, left) and more fMRI-level clustering for those same categories (Fig. 4C, right). A one-way ANOVA indicated a statistically significant difference between the percentage of selective units across different categories (*F* (4, 95) = 27.06, *p* = 5.40 × 10^−15^). Post-hoc Tukey’s honest significant difference (HSD) tests indicated that the percentages of selective units for face and body were significantly higher than for other categories (both *p*s < 0.05). In addition, there was no significant difference in the percentage of units selective for faces and bodies when compared directly (*p* > 0.05). As a measure of the degree to which category-selective regions appear to be clustered in fMRI-like maps, we also defined a “cluster strength” metric for each object category as the sum of the contrast magnitude of the largest cluster in the smoothed maps (see Materials and Methods for details). Using this metric, TDANN model cITs predict that face- and body-selective neurons are more strongly clustered (as measured at the level of reproducible fMRI ROIs) than neurons for other object categories in the test set, *F* (4, 95) = 9.51, *p* = 1.64 × 10^−6^ (Fig. 4C, right). Post-hoc testing (Tukey’s HSD test) confirmed that TDANN cITs predict faces and bodies were significantly more strongly clustered than the other object categories, cars and instruments (both *p*s < 0.05). We found that the TDANN model cIT predicted no significant difference in cluster strength for faces and for bodies when compared directly, *p* > 0.05.

Perhaps not surprisingly, we found that the model’s ability to predict cluster strength for any tested category was strongly, but not perfectly, related to the fraction of model units that were selective for that category (Fig. S7). In other words, the greater the number of category-selective units that the model predicted should be found, the greater the predicted fMRI cluster strength for that category. In particular, most TDANN cIT models predicted high percentage of units selective for faces (median: 7.5%, range: 4.2-21.5% over 20 models) and bodies (median: 10.4%, range: 3.9-17.8%), and low percentages of units selective for instruments (median: 1.8%, range: 0.6-4.4%). While nearly all TDANN models predicted fMRI patches for faces (18 of 20 models) and bodies (20 of 20 models), they rarely predicted model patches for instruments (5 of 20).

To further investigate the existence of selective clusters for a broader range of categories, we tested the selectivity of TDANN model cIT units with the test stimulus set used in (42), which is comprised of gray-scale images of 20 different categories, including faces and bodies. Similar to the findings in (42), we did not find other regions that are as strongly clustered as face patches (Fig. 4D). Significant differences were found among the cluster strength of categories (one-way ANOVA, *F* (19, 380) = 8.18, *p* = 4.01 × 10^−16^). Tukey’s HSD post-hoc tests indicated that face-selective units were significantly more clustered than any other category (all *p*s < 0.05). We note that while the cluster strength for bodies in the model cIT was lower with this stimulus set (Fig. 4D), the body-selective units identified with this stimulus set were localized to the same body sub-regions in Fig. 4A,B.

We note that, unlike primate brains (42, 49), the TDANN layers did not show patches selective for scenes. This may not be surprising as the TDANNs were designed as a model of foveal vision (i.e., the central 8 visual degrees; see Materials and Methods for details) and thus was not intended to model visual areas with peripheral biases in which scene-selectivity is often reported (52–54) (Also see Discussion).

#### Face- and body-selective patches in TDANN model cIT layers often appear adjacent to or overlapping each other

Given the presence of clear face- and body-selective patches in the model cIT layer for nearly all random initializations, what is the spatial relationship between them? In humans and non-human primates, body-selective regions are typically found adjacent to or partially overlapping with face patches (30, 55). We observed a similar spatial relationship between face-selective and body-selective units in the TDANN model cITs. Body-selective TDANN units were often localized near clusters of face-selective units, or intermixed with the face-selective population (Fig. 4A, upper). Quantitative analysis supports the observation that body-selective regions are closer to face-selective regions than expected by chance. We first determined the peak selectivity points for face and body as the center of the most selective 1 mm circular region in each layer (see Materials and Methods for details). The example peak selectivity points are shown as white-colored plus markers for faces and stars for bodies in Fig. 4A and 4B. We then measured the distances between the peak selectivity points of face and body for each cIT layer of all 20 randomly initialized models and found that the mean distance of peak face- and body-selectivity points was smaller than the mean distance between two random points, and was lower than the 98% confidence interval for two random points.

To better compare these results to the fMRI literature in which these patterns were observed in primates, we also investigated the voxel-level overlap in the smoothed maps described above. Visual inspection suggests that the clusters of face-selective voxels were generally adjacent to clusters of body-selective voxels (Fig. 4A, lower), and in many cases these patches partially overlapped one another (Fig. 4B, S6).

#### TDANN model cIT face patches are similar in spatial profile to macaque cIT face patches

We found that both the spatial scale of the model cIT face clusters and the rate at which face-selectivity fell off with distance from the cluster center were similar to precise measures of these parameters in the monkey cIT face patch (aka “middle face patch”; 40). While the spatial constraint applied during TDANN model training should lead to some spatial biasing of similarly responding units, it does not guarantee a precise match between the spatial structure of macaque and model cIT face patches. To more precisely compare the spatial structure of TDANN model cIT and macaque cIT face patches, we computed the spatial profile of the model face patches and compared with the measurements reported in (40). One such measurement is the purity curve, which describes how the fraction of units that are deemed face-selective (by a standardized criterion) falls off as a function of distance from the spatial location with the highest concentration of face-selective units (40). We computed the proportion of face-selective units (purity) in each 1mm x 1mm region of model cIT, then plotted purity as a function of distance from the maximally-selective region (see Materials and Methods for details). We found that these purity curves were similar to those reported from single-unit analyses in the macaque middle face patch (40) in spatial profile, spatial scale, and peak/trough purity values (Fig. 3B, right).

#### Model face-patch units are highly interconnected with face-patch units in other layers

Face patches are sub-regions within each of the three levels of macaque IT (i.e., pIT, cIT, aIT) (41), which is a hierarchical, densely interconnected network that is a continuation of the hierarchical ventral stream (4). Prior work suggests that face patches are preferentially connected with each other across those three levels rather than being equally connected to all neurons in the levels above and below. Specifically, electrical microstimulation of neurons within a single face patch elicits strong activity in other face patches, while microstimulation of neurons located outside of face patches fails to do so (31). To evaluate the interconectivity of the TDANN model cIT and aIT face patches, we mimicked the stimulation paradigm described in (31). Specifically, we simulated microstimulation of neurons both inside and outside of the face patches in model cIT (Fig. 5, S11) and assessed the spatial pattern of microstimulation-evoked responses in model aIT (see Materials Methods). We found that stimulation of model units within each model cIT face patch drove a high level of activity of model units in face patches in the subsequent model layer, model aIT (Fig. 5B). However, stimulation of model cIT locations outside of the face patch did not lead to a significant increase in the activity of face patch units in model aIT (Fig. 5C). A paired t-test across the 20 randomly-initialized models revealed that within-patch model cIT stimulation led to substantially higher activity in the model aIT face patches than stimulation outside of model cIT face patches, *t*(19) = 5.78, *p* < 0.001 (Fig. 5D), providing evidence of interconnectivity of TDANN face patches in successive layers. It is worth stressing that this connectivity was not built into the TDANN models and prior to training, all TDANN models started with each model cIT unit connected to each model aIT unit with random weights (see Materials and Methods).

### Detailed comparison of individual neural units in TDANN model cIT face patches with face neurons in macaque IT

Do the similarities between our models and macaque IT extend beyond topographical phenomenology? To address this question, we compared the responses of “neurons” in model cIT face patches to the responses of actual “face neurons” recorded in macaque IT. We made three types of comparisons: response predictivity, pairwise response correlations over images (a.k.a. signal correlations), and face identity and viewpoint invariance.

#### TDANN model cIT face-patch units predict the neural responses of individual face-selective neurons in macaque

Prior work (1, 36) has shown that non-topographic models trained on general visual recognition lead to model “IT” layers that accurately explain/predict the firing rates in macaque IT. Would the same be true with TDANN IT layers despite the additional spatial constraints imposed during training? The spatial loss function that we applied here to create the new TDANN models during training encourages units in close proximity to have responses that are similar to each other (Fig. 2). One concern from applying this spatial loss function is that since the response properties of all the model IT units will be altered, the resulting TDANN model face neurons might not be as good at explaining the IT face neurons as model neurons from prior studies. We tested how well different populations of model units predict the neuronal activity of macaque face-patch neurons in response to a set of 3,200 images of photorealistic 3D objects drawn from eight natural categories (i.e., animals, boats, cars, chairs, faces, fruits, planes, and tables) with variations in the object position, scale, and pose (47).

We investigated the response profiles of macaque face-selective neurons (*d*′ > 0.5) and TDANN model cIT face-patch face-selective units (*d*′ > 0.66) to different categories (Lower *d*′ threshold values were used in order to ensure sufficient numbers of face neurons). Fig. 6A shows the standardized responses of macaque face neurons (upper) and a subset of model face units (lower) (see Fig. S9 for the full set responses). We observed that the face images elicited stronger responses across both macaque face neurons and model face units relative to the images of non-face objects. We also found that model cIT face-patch units can explain neuronal responses to images of both faces and non-face objects (Fig. 6B), where the Pearson correlation between the profile of category responses for neurons and model units was 0.97.

**Fig. 6.**
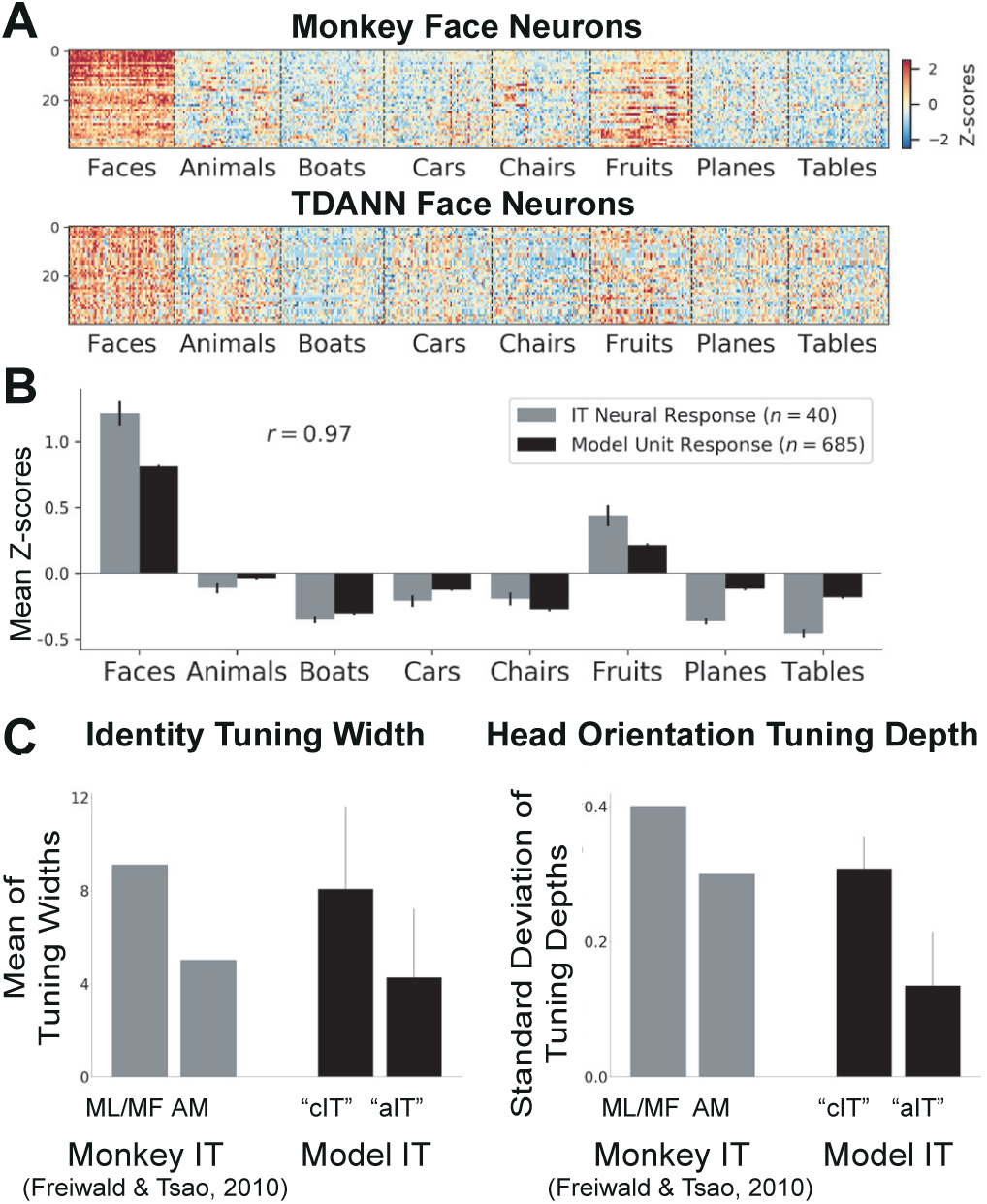
Comparison of monkey IT face neurons and TDANN model IT face neurons. A) Upper panel: Example response profiles over images from eight object categories of a set of face-selective neurons randomly sampled from macaque IT face-selective neurons (*n* = 40, 47). Neural responses were measured as the average firing rate from 70 to 170ms post-image onset (baseline-subtracted using each site’s response to blank gray image as the baseline; see experimental details in (47)). Lower panel: Responses of a randomly-selected sample of face-selective neurons (*d*′ > 0.85) “recorded” from TDANN model cIT face-patches (*n* = 160; see Fig. S9 for the full set of model responses). The responses of each neuron/unit were standardized as z-scores over 512 held-out test images. B) Mean response (over images) within each of the eight object categories for monkey IT face-selective neurons (*n* = 40) and face-selective neurons from TDANN model cIT face patches (*n* = 685). Error bars indicate the SEM over neurons or model units. C) Facial identity and viewpoint tuning in macaque and TDANN model cIT/aIT face patches, following the conventions of (19). Left panel: Mean (over neurons) identity tuning widths for neurons sampled from monkey face patches (left; ML/MF and AM) and neurons sampled from TDANN face patches (right; model cIT and model aIT). Smaller tuning half-widths indicate greater identity selectivity. Right panel: Mean (over neurons) of standard deviations of head orientation tuning depth. Tuning depths close to 0 (and thus smaller standard deviations of tuning depths) indicate greater viewpoint invariance. Error bars indicate the SEM over model units. The values for macaque face patches were reproduced from the reported values in (19).

We also tested the ability of TDANN model cIT face-patch units (*d*′ > 0.85) to predict responses of face-selective neurons in macaque IT (*d*′ > 0.65). (47) notes that most of these face-selective neurons are likely belonging to the IT face patches. Following the procedure in (36), we used a simple partial least squares (PLS) regression to predict the activity of each recorded macaque face-selective neuron from a linear combination of model face-patch units. The predictivity scores were then measured as the noise-corrected Pearson correlation between the actual and predicted responses of each neuron on a held-out set of images. We found that model cIT face-patch units explained the activity of macaque face-selective neurons well. The median correlation between predicted activity and actual activity was 0.623 *±* 0.016 (mean *±* std) across 20 models with different random initializations. Fig. S10A shows an example distribution of the neural predictivity scores for one exemplar model.

We additionally tested the ability of model cIT face-patch units to explain face-patch neurons in macaque IT using the data collected in (40). Unlike (47), the neurons recorded in (40) are specifically localized within or nearby face patches. We used a subset of the neural data that was collected: 250 images spanning macaque faces, human faces, scenes, objects, and body parts. We used cross-validated Ridge regression to predict the activity of each recorded macaque face-patch neuron from a sparse linear combination of model face-patch units. The median predictivity score (noise-corrected) was 0.645 *±* 0.032 (mean *±* std) across 20 models with different random initializations. In other words, about 42 percent of the explainable visually-driven response variance of face neurons is explained (predicted) by the TDANN model cIT (this is comparable to other (non-topographic) deep ANN models on similar tests, unpublished). However, it should be noted that these results likely underestimate TDANN’s ability to explain face-patch neuron responses due to the low number of images that were available to map the model units to the recorded neurons (see Materials and Methods for more details). Nevertheless, the model cIT face-patch units could fit the activity of macaque face-patch neurons well overall.

These results suggest that the addition of a spatial loss term during model training does not prevent the formation of face-selective model units in ANN models of the ventral stream and it does not degrade the functional similarity of these units to face-selective macaque neurons. Does it degrade the ability of the total distribution of model units to predict the activity of a broad population of IT neurons?

To answer this question, we next investigated the ability of all model units to predict IT macaque neurons regardless of face selectivity (Fig. S10B). We compared the predictivity of model units in the model cIT to units from model aIT, as well as earlier model layers whose activity might more closely reflect that of early and intermediate visual areas. A one-way ANOVA revealed a significant difference between the quality of fits from different TDANN model layers (green line in Fig. S10B), *F* (6, 133) = 23788.7, *p* < 1 × 10^−198^, and post-hoc tests indicated that fit correlations were higher in model cIT than in all earlier layers (all *p*s < 0.05). Furthermore, fit correlations from model aIT were significantly higher than fits from all layers earlier than CONV5 (all *p*s < 0.05).

We also compared layer-wise fits between the TDANN and non-topographic control ANN models. We conducted a 2-way ANOVA (model type × layer) which indicated a main effect of model type, *F* (1, 126) = 56.7, *p* = 8.47 × 10^−12^ and a significant interaction between layer and model type, *F* (6, 126) = 39.4, *p* = 1.11 × 10^−26^. A post-hoc test indicated that some of the TDANN layers are better in explaining the neural responses than the control model layers (CONV3, CONV5, and model aIT; all *p*s < 0.05). Yet, overall there was no significant difference between other layers of TDANN and control models (*p* < 0.05).

#### The response similarity of model cIT face neurons is similar to the response similarity of monkey IT neurons

Having established that the TDANN models explain neuronal responses at a level consistent with prior ANN models, we further tested if the additional spatial constraint resulted in model units that were more (or less) correlated with each other than the actual pairs of IT face-selective neurons from the same face patch. We computed the mean pairwise correlation between responses from model units in TDANN cIT face patches (1 patch from each of the 20 models) and compared to the mean pairwise correlation of neurons in 7 macaque face patches from 2 different monkeys. For both model units and macaque neurons, correlations were computed over a set of natural images used in prior studies (40). The mean pairwise correlation of macaque face-selective neurons from the same patch (*r* = 0.59 *±* .04) was not significantly different from the mean pairwise correlations of face-selective model units from the same patch (*r* = 0.54 *±* .01), independent samples t-test *t*(25) = 0.89, *p* = 0.41, suggesting that the TDANN model face-patch units exhibit similar correlation structure to face-patch neurons in macaque IT.

#### Identity selectivity and viewpoint invariance increases in successive TDANN model layers

In the macaque ventral stream, invariance to object viewpoint in neural representation that convey visual information increases from posterior to anterior IT (19), consistent with the cortical hierarchical progression of invariance for general object representation (8). For coding of face identity in particular, results suggest that neurons in posterior IT face patches respond to any facial identity at a given viewpoint, neurons in anterior face patches respond to a given identity regardless of viewpoint, and intermediate regions respond similarly to mirror-symmetric viewpoints (19). We evaluated the face viewpoint invariance of the TDANN model cIT and model aIT face-patch units by presenting images of 25 different faces at 8 viewpoints each (Fig. S8A, from 19) and computing identity and viewpoint tuning in face patches.

Following the stimuli and the procedure in (19), identity tuning width and head orientation tuning depth were computed for each face patch unit across all models (see Materials and Methods for details). Here, smaller identity tuning width corresponds to higher selectivity to identities, while smaller head orientation tuning depth means greater viewpoint invariance. We observed an increasing identity selectivity (Fig. 6C, left) and an increasing viewpoint invariance (Fig. 6C, right) from TDANN model cIT to model aIT. Indeed, TDANN model aIT identity selectivity was significantly greater than model cIT (*p* << 0.001, Mann-Whitney U test, Fig. S8B), and TDANN model aIT viewpoint invariance was significantly greater than model cIT (*p* << 0.001, Mann-Whitney U test, Fig. S8C). These trends are consistent with those reported for the macaque face path system (19). In summary, the previously reported trend of increasing identity selectivity and viewpoint invariance from ML/MF (located in cIT) to AM (located in aIT) was observed between TDANN model cIT faces patches and model aIT face patches.

### Dependence on visual diet

In this work, each TDANN model was trained (i.e., evolved or developed; see Discussion) with naturalistic images from the ImageNet (ILSVRC-2012; 45) and Labeled Faces in the Wild (LFW; 46) datasets. The ImageNet dataset used in the ILSVRC-2012 competition is a large-scale image dataset comprised of 1000 categories built upon the WordNet (56) synset tree. While its large scale and diversity has led to substantial advancements in computer vision, it is not clear whether the ImageNet dataset reflects the actual visual experience “diet” of primates. For instance, 87 of the ImageNet classes are different types of vehicles, whereas there are very few classes of humans or other primates. Even for humans who, unlike other primates, may be exposed to some of those vehicle types, it is unlikely that they experience more vehicles than faces or other humans, especially during evolution and early visual development.

Recent neuroscience findings (57, 58) suggest that the early visual experience of primates affects the development of categorical representations in IT. Extending this idea to models, we asked if and how the representations in the TDANN model cITs and aITs depend on their visual experience by training the networks using datasets with different categorical distributions. Here, we define the “visual diet” of each TDANN model as the combination of the set of images used during training and the corresponding set of recognition tasks associated (i.e., the set of ground truth image labels). For this experiment, three additional modified training sets were constructed from the ImageNet and LFW datasets by manipulating the frequency of face- and vehicle-related categories (Fig. 7A) in the training set. We trained 10 new TDANN models for each visual diet and compared both the percentages of category-selective units and the cluster strength of category-selective units in model cIT and aIT with the same measures we used in the baseline TDANN models (described in Fig. 4).

**Fig. 7.**
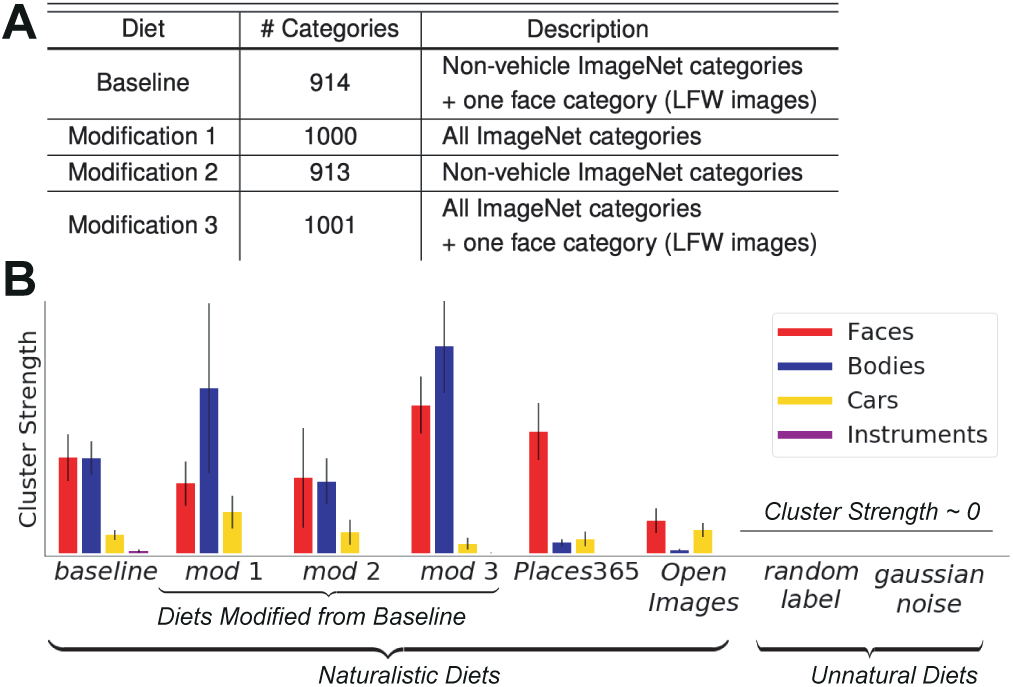
The effect of visual experience (training) diet on the cluster strength of category-selectivity (*t* > 10) in TDANN model cITs. In panel B, the first six diets are naturalistic: the baseline diet, three diets modified from the baseline diet (*modification 1-3*; subsets of ImageNet (45) and LFW (46) as shown in panel A), and two entirely non-overlapping sets of images: Places365 (natural scenes; 59), and Open Images (naturalistic images; 60), from left to right. The two rightmost diets are highly unnatural: ImageNet (45) images with randomly shuffled labels (*random label*) and random Gaussian noise images (*gaussian noise*; see Materials and Methods for details). The plot shows the mean cluster strength (mean over all TDANN models trained with the indicated diet) for faces (red), bodies (blue), cars (yellow), and instruments (purple). For each category, the cluster strength was measured using the test images from (25), using the same (∼fMRI) cluster strength metric as in Fig. 4C. Error bars indicate the SEM.

We found that in model cIT, there was no significant difference in either the predicted percentage of units selective for faces, bodies, cars, and instruments, or the predicted cluster strength for faces, bodies, cars, and instruments (all *p*s > 0.2; the first four columns in Fig. S12A and 7B, respectively). However, relative to the baseline diet, the aIT level of models trained on standard ImageNet (modified diet 1) had a 12.5% smaller percentage of face-selective units (*t*(28) = 2.01, *p* = 0.05), a 9.9% larger percentage of car-selective units (*t*(28) = −4.25, *p* = 0.0002), and no change in percentage of body-selective units (*p* > 0.4) (the first two columns in Fig. S12B). These differences in the proportion of units selective for each category was accompanied by higher cluster strength for face-selective units (*t*(28) = 2.46, *p* = 0.02), a lower cluster strength for car-selective units (*t*(28) = −3.53, *p* = 0.001), and no change in the cluster strength of body-selective units (*p* > 0.5), as shown in Fig. S12C (first two columns). These results suggest that, for TDANN models, the training diet has its largest effect at the very highest (“output”) layers (closest to the behavioral task), but they also show that the core phenomena of clusters of face neurons (and body neurons) in the more “middle” model layers (approximating cIT), are relatively robust to (these) changes in diet.

We also found that the other hallmarks of face processing were robustly found in TDANN models regardless of the training diets: the models’ ability to explain the response profiles of individual macaque face neurons, the inter-connectivity of face patches, and the increase of viewpoint invariance in successive layers were unaffected by the choice of visual diet (Fig. S13).

Given the robustness of face-patch properties across different categorical distributions, we next asked whether the model predicts that face patches will appear with even more drastic changes in the visual diet. In this experiment, we trained new TDANN models on four additional diets: 1. Places365 (59), 2. Open Images (60) in Tencent ML-Images (61) format, 3. random Gaussian noise images, and 4. ImageNet images with randomly shuffled labels (see Materials and Methods for details). We trained ten new models on each dataset. The first two diets are naturalistic images, but are intended to further test robustness. The latter two diets are both highly unnatural in different ways, and were included to test whether the face patches would appear under nearly any visual task training regime. Using the same images and measures of face cluster strength as above (the rightmost four columns in Fig. 7B and S12A), we found that face clusters still appeared in models developed on the two natural diets (1 and 2). On the other hand, we found that training the models with randomly labeled naturalistic images or random noise images generally destroyed the appearance of the face network in the model (see below for an additional robustness analysis), suggesting the importance of ecological visual experience.

### Robustness of results to operational definition of face selectivity

As a final test of robustness, we asked how strongly the model’s predictions of face neuron appearance and spatial clustering depend on the images sets that are used to operationally define face neurons in the first place (i.e. the positive (face) and negative (non-face) test images). In particular, while all of the results presented above used standard operational test images from the neuroscience literature to facilitate direct comparison with neural and fMRI results, we noticed that measures of model face selectivity had some dependence on the operational definition (e.g. Fig. 3A).

To explore the impact of this, we tested how the face selectivity of each TDANN model’s units depended on the test images. Specifically, for each TDANN model, we computed the spatial pattern of face selectivity (d’) values over all TDANN cIT model units using each of the four test sets (25, 40–42), and we measured the reliability of this pattern of face selectivity by computing the correlation of the d’ values over all units (see Supplementary Information for details). Fig. S14B shows the average reliability scores of the TDANN models trained with different diet as the mean pair-wise correlation of d’ values. We observed that the models trained with naturalistic datasets (leftmost six columns) were relatively highly correlated (mean: 0.42), but not perfectly correlated, suggesting some dependence on the test sets. We do not know of any neuroscience study that makes similar comparisons across test sets, so we cannot ask if this level of (non-perfect) robustness to the image test set is also found in individual brains. As a reference, we also found that models trained on very unnatural diets (rightmost two columns) showed very unreliable patterns of face selectivity (mean: 0.06), indicating that, while they may occasionally appear to demonstrate “face selectivity” under one operational definition, it is clearly not face selectivity as it does not hold up under changes in the details of the test set. Interestingly, we also found that training with naturalistic images assigned to random labels showed slightly more reliable spatial patterns of face selectivity than the random noise training. This suggests that while learning the semantic meaning of visuals is important in the emergence of face patches, simply experiencing naturalistic images may also contribute to the development of categorical representations.

## Discussion

In this work, we tested the hypothesis that the neurobiological hallmarks of specialized face processing in high level ventral visual cortex might be an emergent consequence of the ventral visual stream evolving and/or developing to solve general visual recognition while also minimizing wiring costs. To test this hypothesis, we built new artificial neural network models that implement those two goals. Specifically, we assumed that the evolutionary/developmental pressure to minimize wiring cost could be approximated by encouraging physically nearby neurons to have similar response profiles (see discussion below). Thus, to test the overall hypothesis, we trained neural networks to solve real-world visual object categorization tasks (as in prior work; 36) while *also* striving for a solution in which neurons with similar response profiles are physically close to each other (referred to as a “spatial correlation” cost). We do not assume that optimization procedure to minimize these costs (gradient descent training) is emulating the same mechanisms of evolution and development – only that it provides a way to apply the two types of fitness pressures on a network of similar architecture to the ventral stream and observe what internal representations it tends to develop.

We succeeded in meeting the engineering challenges of applying these two pressures, which resulted in neural network models in which the highest layers of those models had artificial neurons arranged in two physical tissue sheets (cIT) and (aIT) and their weights are optimized such that the nearby neurons show similar response profile, and also performed well on real world object categorization. We refer to these models as topographic deep artificial neural networks (TDANNs). We then probed these TDANN models in a manner directly analogous to neuroscience experiments in humans and non-human primates to characterize and systematically compare the empirical neurobiological hallmarks of face processing. We mostly focused on monkey IT cortex (specifically central and anterior IT; cIT and aIT) and referenced human data when comparing fMRI maps. Since a similar network of face patches are found in human ventral temporal cortex, we expect the qualitative phenomena will be similar and thus relevant to both species.

Our results reveal that these TDANNs reproduce many of the neurobiological hallmarks of face processing in primate higher visual cortex. Most notably, we observed clustering of face-selective units into “face patches”, that the purity and spatial extent of those face patches was very similar to that measured in monkeys, enriched connectivity between face patches in different model layers, and the emergence of viewpoint invariance in successively higher face patches. Furthermore, we observed that neural units in TDANN model face patches predicted the responses of individual macaque face-patch neurons, and these predictions were comparably accurate to those of earlier neural network models of the ventral visual stream (1, 12, 36).

To us, the most salient finding was the reliable emergence of face (and body) patches, even though faces were not explicitly treated as special and the spatial correlation rule was not tuned in any way to require those structures to emerge. Indeed, we tested other standard categories of objects that, a priori, should have been equally likely to have physical clusters of neurons. However, we rarely observed the emergence of patches for those categories, just as prior fMRI work in humans has never reported patches for these same categories, despite intensive efforts to do so (42, 50, 51). This salient finding of face (and body) patches is even more striking when we find that their emergence is highly robust to the choice of training diet (experienced images and tasks). However, the TDANN models predict that highly unnatural visual diets prevent the formation of these structures (see below).

Taken together, these results show that TDANN models readily explain the main reported empirical hallmarks of the IT face processing network, even at a quantitative level. Thus, those empirical hallmarks do not by themselves imply developmental mechanisms that are orchestrated specifically to create a face network. Instead, they suggest the hypothesis that some relatively simple developmental rules might be sufficient, even over a broad range of natural visual experience (Fig. 7, S12 and S14). An important caveat is that, because we do not assume that the training of TDANN models is a mechanistic model of development, our results do not by themselves rule out the possibility that evolution has effectively incorporated wiring cost pressure (pressure that is approximated by the training of the TDANN) into the genome which then in turn orchestrates a more detailed construction of the ventral stream, including face patches. Indeed, empirical evidence on this question is conflicting (cf. 57 versus 62, 63 and 64).

Regardless of the above, the general inference we draw from this work is that the functional organization of the ventral stream as a whole might be understood as resulting from the need to perform behaviorally important inferences of latent visual image content (esp. object category) while also minimizing wiring costs over evolution and/or post-natal development.

### Spatial correlation rule as a proxy of wiring cost minimization

The overarching hypothesis we tested – cortical topographic organization emerges from solving general visual recognition in the face of neural wiring costs – is not conceptually new (65–67). But, to our knowledge it has not been previously tested in higher visual cortex because our field did not have neural network models that could reasonably explain and predict the response patterns of individual IT neurons, let alone the specific physical arrangement of face-related neurons in the IT tissue. However, the advent of reasonably accurate, image- predictive artificial neural network models of the entire ventral visual stream (1, 12, 36, 68–70) suggested this hypothesis could now be tested. Our work here builds upon those models and an overall performance optimization approach (1).

The spatial correlation cost that we developed and required each TDANN model to minimize is inspired by and consistent with the hypothesis that the brain should perform its necessary computations with minimal axonal wiring cost. This framework posits that physical and metabolic constraints have, over evolution and/or development, favored a spatial organization of neurons in each cortical area that minimizes the length of axons and extent of dendritic arbors (71, 72). To implement this underlying motivation in an engineered model, we did not directly compute the wiring length cost of each possible arrangement of all the neurons in an area, but we instead used an efficient proxy. Specifically, we assumed that wiring cost minimization should tend to lead to physical arrangements of neurons in which neurons with similar response profiles are near to one another, so we could optimize for that goal. The reasoning here is that the circuit drivers of those response profiles (as implemented through axonal connections) would tend to be reduced in this physical arrangement (relative to a physical arrangement in which, for example, neurons are randomly arranged in the cortical tissue) (33, 34). Using basic assumptions about the wiring cost needed to implement any given arrangement of neurons, we do find that TDANN cIT layer (optimized with the spatial correlation cost) do indeed have lower wiring costs than random spatial arrangements of neurons from the same base architecture (AlexNet) trained on the same set of categorization tasks (see Supplementary Information for details). However, an important goal of future work is to develop and implement more direct methods of wiring minimization in deep ANN models, and to implement those methods at all levels of the model ventral stream (not just within the upper “IT” layers as we have done here). While we fully expect such models to reproduce the basic findings described here, they could be even more accurate models of the entire ventral stream, and would allow a more direct exploration of the metabolic and physical costs that might have been most important to shaping its functional organization.

### Intuition about neural clustering

We found that the proportion of selective units that emerged for each category, was highly correlated with the clustering strength of those units on the cortical surface and thus with the likelihood that mm-to-cm scale measurements such as fMRI and optical imaging would detect reliable signals for those categories (Fig. S7). That is, the TDANN models suggest that it is suboptimal to form clusters for categories with low numbers of selective units, such as tools or flowers (Fig. 4). This may be intuitively seen as a natural consequence of the spatial cost function the TDANN aims to satisfy, and it remains to be tested if it holds up under other related cost functions (such as more direct measures of wiring cost or energetics). But, even if the spatial clustering itself seems intuitive, that intuition does not at all explain what determine the fractions of units that are selective for each category.

We also observe the fraction of selective units, the clustering strength, and the precise spatial location of the model face patches (e.g. Fig. 3A), are partially dependent on the operational definition of selectivity used to probe the neural network, i.e. the image test sets. For instance, TDANNs robustly predict the face selectivity across different test sets, but they predict some differences for body selectivity (e.g. lower clustering strength for bodies with some test sets; see Fig. 4C,D). To our knowledge this has not been systematically tested in the ventral stream, and further neuroscience investigation is needed to identify if the subtle, but reliable variations seen in the TDANN models are in line with those in individual brains. Indeed, this is one of several areas where the TDANN models motivate future experimental work that would drive further model development.

Our manipulation of the training diet, along with similar experiments in macaque (57), suggests that visual experience is one important influence on the proportion of units selective for a given category. We are unable, however, to rule out other possibilities, including factors related to the distribution of visual features that define a category or yet-to-be-discovered genetic mechanisms that guide the distribution of category-selective units (see below).

### Dependence on visual diet

We observed that the distribution of images experienced by the TDANN model and the specific set of tasks it was asked to perform on those images (together referred to as the model’s visual “diet”) affects the final “adult” set of neurons and their physical arrangement within the model cortical tissue. For example, the removal of the face detection task and the associated images reduced the number of face-selective units, and it reduced, but did not eliminate, model face patches. Similarly, the inclusion of ImageNet’s vehicle-related categories in the diet increased the car-selectivity, but this effect was most pronounced in the last hidden neural layer (fc7, model aIT) (Fig. S12). We also observed that the TDANN layers showed low selectivity to scenes that were not included as an explicit category during training. Although some training images may contain scenes as backgrounds, the general visual recognition task the models were trained on did not include “scenes” as a target label.

Yet, interestingly, we found that models trained with the standard ImageNet category diet – which seems very non- natural from a evolutionary task perspective – already contain large numbers of face- and body-selective units and the clustering of such units (relative to other categories, such as cars and instruments). Indeed, we noted that even with no face detection or face discrimination task in the training diet, most TDANN models showed face-selective neurons in large enough numbers to result in face patches. One interpretation of these findings is that the ability to distinguish face shapes, for example, may come at least partly for free from the high level shape features that implicitly result from learning how to solve a general object-categorization tasks. Another, similar interpretation is that because the naturalistic image sets used in training all contained at least some images with faces, and the presence and properties of those faces were likely diagnostic for some trained categories, the existence of face patches could reflect the implicit benefit of detecting and/or discriminating faces for categorizing objects and scenes. Further, our results also suggest that the strength of face clustering positively depends on the amount of faces in the visual diet and/or tasks associated with discriminating faces (we did not attempt to disentangle those two factors here as that is non-trivial and is the focus of future work).

Given the robust emergence of face (and body) patches with a range of diets (above), we began to wonder about the minimal requirements needed to see their emergence. While we have not exhaustively tested all the possible variables, we did find that diets that are strongly unnatural in terms of image statistics (Gaussian noise images with fixed labels) or are strongly unnatural in terms of tasks (random labels assigned to natural images) do not reliably produce face patches in the TDANN model (Fig. 7, S14).

If one assumes that the TDANN training is very crudely approximating mostly post-natal development, then the diet results (above) are at least partially consistent with the finding that non-human primates deprived of face experience tend to have weak to non-detectable face patches, and that the patchiness of the IT functional organization depends on tasks learned during post-natal development (57). However, in our opinion, the modeling work is not yet sufficiently advanced to appropriately engage on these questions, as the model training is highly non-biological in that it is fully supervised.

### Directions for future work

The modeling results presented here naturally lead to a number of questions for future work: what is the distribution of different categories that best reflects the true visual experience of primates? Can we build a more ecologically balanced set by tuning the distribution based on the similarity between the brain’s and models’ representations? Here, we explored a limited range of training diets, as a full exploration of all possible training diets is beyond the scope of this work. Future work will be needed to determine how visual diet affects the development of categorical representations and the formation of different patches. Studies of that form will ultimately require: constructing datasets that are closer to the primate visual experience (73), unsupervised and or self-supervised model learning methods (74), and comparison with data from primates at different stages of post-natal development (75) and different types of visual experience (57). This would allow future modeling work to begin to separate the mechanisms of evolution (i.e. defining a model’s “birth state”) from the mechanisms that are work during post-natal development and adult learning.

Another future engineering direction is to develop ANN models that apply spatial/wiring costs at all levels of the ventral stream – not just within the IT levels as we have done here. For current deep ANN models, engineering challenges must be overcome, but progress is underway (76). And a related direction is to switch from the baseline deep ANN architecture used here (AlexNet) to other more advanced ANN architectures that better approximate the recurrent circuitry both within and beyond the ventral stream (77–79).

On the neuroscience side, the TDANN models already motivate several new lines of experimental work. For example, more precise measurements of the dependence of the face and body patches on the specifics of the image tests sets (outlined above). Even more interestingly, the TDANN models can be used to predict sets of images that should best activate or deactivate tissue regions of IT that lie outside the known categories. Indeed, older work (80, 81) and more recent work (82) suggest that, even at the level of fMRI, these other regions of IT are indeed selective over some image set contrasts. But the TDANN models likely offer even more precise predictions.

Clearly much remains to be done. Even though the TDANN models reveal how the hallmark neural phenomena of face processing might be explained, these models are certainly incomplete in many aspects. However, they already suggest new experiments that will in turn lead to new models that will advance our understanding. Indeed, our field is only at the cusp of applying such image-predictive models to explore, predict, test, and ultimately understand these and other natural phenomena in the visual system. This iterative loop between engineered models and experiments holds the promise to elucidate conceptual hypotheses and resolve long standing debates, and this approach will accelerate as our field adopts and deploys engineered models even more advanced than those we deployed here.

## Materials and Methods

### Spatial correlation loss function

The spatial correlation rule was implemented as a proxy of wiring cost minimization within each of two layers of the ANN architecture (Fig. 2). Here, the underlying assumption is that the pressure to minimize wiring cost can be approximated as a pressure to have neurons with similar response profiles to be adjacent. Inspired by that, we derived the spatial loss function by considering pairwise similarities of model unit responses in each layer as a function of their physical distance within the tissue map of that layer. This heuristic is consistent with observations that pairwise neuronal correlations in visual cortex decrease as a function of distance on the cortical surface (83, 84). The specific loss function we used was parameterized to approximate the actual IT response correlation spatial profile that were obtained from the neural recordings (47) in macaque IT cortex (Fig. 2):

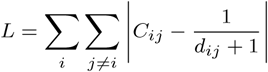

where *C*_*ij*_ is the response profile correlation between the model units *i* and *j*, and *d*_*ij*_ is their cortical distance in millimeters in the tissue map.

### Assignment of unit tissue positions and model training

The topographic deep artificial neural networks (TDANNs) in this work are adaptations of the AlexNet (2) base architecture: five convolutional layers with max-pooling nonlinearities after layers CONV1, CONV2, and CONV5, followed by two fully-connected layers (FC6 and FC7; Fig. 1). In our TDANN implementation, model units in layer FC6 and layer FC7 are assigned initial, random spatial positions in a 10mm x 10mm two-dimensional map, simulating the approximate amount of cortical tissue in each level of IT devoted to the central ∼8 degrees. Separate, independent model maps were maintained for each layer. Once training began, the model neurons all remained physically fixed in the tissue, but the internal synaptic weights to each neuron were iteratively adjusted to optimize the model image categorization performance (see below) and to minimize the spatial loss (above). The overall spatial loss in each layer was set to be ∼0.25 of the overall categorization loss.

Each network was trained using stochastic gradient descent with a batch size of 256 images, momentum of 0.9 and weight decay of 0.0005. The learning rate was initialized as 0.01 and decreased twice by a factor of 10 throughout 100 epochs of training. Twenty TDANN networks were initially trained with random parameter initializations. New networks were later trained to explore the effects of visual diet (see below). As a non-topographic control, ten networks with the same baseline AlexNet architecture were trained with the exactly same procedure but without the additional spatial loss. Unless otherwise noted, all statistics were computed using data from all models.

### Training dataset

The baseline TDANN models were trained with a set of 914 visual categories from a naturalistic image dataset collected from the ImageNet dataset used in the ILSVRC-2012 competition (45) and the Labeled Faces in the Wild (LFW) dataset (46). To construct a training set a bit closer to non-human primate visual experience, our initial baseline training diet used 913 non-vehicle-related categories (∼1.17 million images) from the ImageNet dataset plus one ‘face’ category (9,500 LFW images from the LFW image set). Thus, the general visual categorization training was for 914 equally weighted categories, one of which was a ‘face’ category. No face-identification training was performed.

To investigate the effect of visual training diet on category selectivity in TDANN models, we trained new TDANN models with one of the three additional training diets (Fig. 7A). Modified diet 1: all ImageNet categories (1000 categories, ∼1.2 million images). Modified diet 2: Identical to baseline diet except no extra LFW face category (913 categories, ∼1.17 million images). Modified diet 3: all ImageNet categories plus the one LFW ‘face’ category (1001 categories, ∼1.18 million images). We trained ten new models on each of these diets.

In addition, we trained new TDANN models with two additional naturalistic training datasets: Places365 (59) and Open Images (60). The Places365 dataset consists of 1.8 million naturalistic scene-centric images assigned to 365 categories. For Open Images, we used 1.2 million images assigned to 1,134 categories, subsampled from the Open Images images which were collected as Tencent ML- Images (61) and re-labeled to match the ImageNet labels. We also constructed two additional datasets that are strongly unnatural in terms of image statistics (*gaussian noise*; Gaussian noise images with fixed labels) or tasks (*random label*; random labels assigned to natural images). For *gaussian noise*, 1.2 million Gaussian noise images were generated from random normal distribution (0,1). The pixel values were clipped to (−1, 1), then rescaled to (0, 255) in order to represent the RGB values. We also trained new models with ImageNet images with randomly shuffled labels, which destroys the semantic relationships between the images and the labels. To maintain the distribution of the labels in the original dataset, the labels of the entire training set were randomly shuffled, and then assigned to the images.

### Visual test stimuli and neural data

In this work, six different visual test sets from (19, 25, 40–42, 47) were used. Each stimulus set contains gray-scale or colored images from face and non-face categories, such as objects and bodies. For neural predictivity analyses, subsets of stimuli and neural data from (47) and (40) were used. For neural data, the multi-unit response to each images was recorded from two macaque subjects as the average firing rate in a 70-170 ms (47) or 60-160 ms (40) window after stimulus onset. All responses were then baseline subtracted using the background response to blank gray images (47) or the average firing rate in a 0-50 ms window after stimulus onset (40). More details for neural data can be found in (47) and (40).

### Simulated fMRI maps

To simulate maps of the same spatial resolution of high-resolution fMRI, we spatially smoothed the activation maps with a Gaussian kernel of full width at half maximum 1mm (Fig. S5). The “voxel” resolution of the smoothed maps was 0.5mm x 0.5mm. Then, the selectivity metric was computed on a per-voxel basis to obtain the voxel selectivity maps.

### Selectivity metrics

Following the procedure in (40), the model units were considered category-selective if they preferentially responded to a given category over other control categories. The control categories were either non-face objects or all other categories in the test set, depending on the analysis. Category selectivity of model units was primarily measured as *d*′. Selectivity to images of faces versus objects, for example, was computed as:

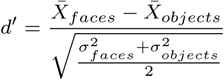

where 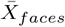 represents the mean response of a unit to all images of face objects, 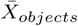 represents the mean response to all images of non-face objects, 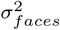 is the variance of the responses to face objects, and 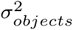 is the variance of the responses to non-face objects.

For a fair comparison between the macaque neural data and model units, we chose the category selectivity criterion (*d*′ > 0.65) used in (40) and translated the threshold value into the model space by approximating the known levels of neural trial-by-trial variability (a.k.a. “noise”). More specifically, we added independent Poisson noise to each model unit responses and computed *d*′ values for both original model responses and noise-simulated model responses. Then, the relationships between *d*′ values of original and noise-simulated responses were fitted to a linear function, which was then used to compute the *d*′ threshold value for the models (0.85) that corresponds to the value from the neural data with noise (0.65). The threshold of *d*′ > 0.65 is approximately equivalent to an *FSI* > 1*/*3, where the responses to faces are at least twice as strong as response to non-face objects, and was derived from macaque middle face patches by (40). For response profile analysis (Fig. 6, S9), the lower threshold values were used for both neurons (*d*′ > 0.5) and models (*d*′ > 0.66) to ensure sufficient numbers of face neurons.

The selectivity for simulated voxels was computed as:

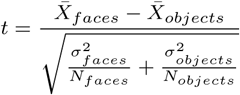

where *N*_*faces*_ is the number of face images and *N*_*objects*_ is the number of object images. The selectivity criterion of voxels was *t* > 10, *p* < 0.001.

### Model patch definitions

To define model face or body patches, we computed the mean *d*′ selectivity of all units in circular regions with 1 mm radius on each map. The peak selectivity point for each category was determined as the center of the most selective circular region. Then, the units within 3 mm distance from the defined peak selectivity point were used for face- or body-patch unit analyses. For neural predictivity analyses, only the face-selective units as defined by 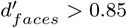 criterion were sampled. For patch connectivity analyses, we used all voxels that are category-selective (*t* > 10, *p* < 0.001).

### Random peak selectivity points

To compare the distances between peak face- and body-selectivity points to random distances, we draw 10,000 pairs of random peak selectivity points. To match the condition with that of actual peak category-selectivity points (see details in the above section), each of random peak selectivity points was drawn from a uniform random distribution over [0 mm, 8 mm] such that each peak point can represent a valid circular region with 1mm radius.

### Measurements of purity

Following the procedure in (40), purity was measured by counting the fraction of face-selective units in every non-overlapping 1mm x 1mm region of the simulated cortical sheet. The region with the highest purity was considered the face patch center, and in every other 1mm x 1mm region, the average distance to the center was computed. To investigate the purity falloff curve, all units at each radial distance from the center were binned at 1 mm resolution. Fig. 3B (right, green curve) shows the mean purity over 20 models in each bin, with an error band representing the SEM across models.

### Measurements of cluster strength

To investigate which category-selective units form spatial clusters, “cluster strength” was computed as a measure of how much category-selective regions appear to be clustered in simulated fMRI maps (Fig. S5). For each layer, we first computed the selectivity of voxels by contrasting each category against all other remaining categories, using one half of the images and corresponding model responses. Then, the region of interest was defined as the largest contiguous category-selective voxels (*t* > 10, *p* < 0.001). For each ROI, cluster strength was estimated as the sum of contrast response intensity (mean response to the category - mean response to other categories) using the other half of the images and corresponding model responses. Fig. 4C (right) and 4D show the mean cluster strength over 20 models, with error bands representing SEM. For this analysis, we used a subset of images from (25) or (42).

### Patch connectivity

For patch connectivity experiments, we first identified the face-selective regions and their center (orange lines and red plus marker, respectively, in Fig. 5) using the half of the stimulus images from (40) for both model cIT and aIT. The center of each face patch was determined as the center of the most selective 1 mm circular region in each model area. The face-selective regions were defined as all category-selective voxels. Then, the stimulation site in model cIT was defined to be a 4 mm circular region centered at the identified face-patch center for inside-patch stimulation. For outside-patch stimulation, the center of the stimulation region was randomly chosen from outside of any identified face patches in the map. The stimulation of model cIT units was simulated by setting the activations of each unit as the product of its maximum activation values over the other half of images and a spatial decay factor sampled from the standard normal distribution centered on the point of stimulation. The units outside of the 4mm circular region were silenced. The observed response in model aIT was estimated as the corresponding activations of model aIT units in response to the simulated activations in model aIT. Fig. 5B,C and S11 show the resulting maps.

### Model prediction of neural responses

To compare the response profiles of face neurons and model face units, we first identified face-selective neurons (*n* = 40, *d*′ > 0.5) from all macaque IT neurons recorded in (47) and face-selective model units (*n* = 685, *d*′ > 0.66) inside the face patches in all 20 TDANN model cIT layers. The selectivity of neurons and model units were computed using 640 images (a subset of low and medium variation sets in 47). The responses of identified face neurons and units were measured using the held-out test images (501 from low variation set).

In addition, we measured the neural predictivity of model units to evaluate how well the responses of model units to given images can explain the responses of macaque IT neurons, using neural data from (47). Linear regression was used to map from model units to each of neural sites with five of 60%/40% train/test splits. We used PLS regression procedure with 25 retained principal components. For each neural site, the predictivity score was estimated as Pearson correlation coefficient between actual and predicted responses. Then, the predictivity score was noise-corrected by normalizing the correlation coefficient by each site’s trial-by-trial variability. The variability for each neural site was computed as Spearman-Brown corrected split-half self-consistency over image presentation repetitions. This procedure was conducted independently for each randomly-initialized model. We also measured the neural predictivity on responses of macaque IT face-patch neurons from (40). In this case, Ridge regression procedure was used with 20 splits of cross validation using 50%/50% train/test splits and 20 retained principal components. Here, we used Ridge regression instead of PLS regression as the data is small and sparse. We note that this particular neural data used a very small set of images (250) that include faces, bodies, objects, and places. Furthermore, not all images were shown to all neurons due to experimental constraints. Due to the small size and sparseness of data, the measured predictivity scores may underestimate the model ability to explain neural responses. For Fig. S10A, only the face-selective units (*d*′ > 0.85) in TDANN model cIT face-patch were used to fit the face-selective neurons (*n* = 34, *d*′ > 0.65). For evaluating the effect of additional spatial constraints on predictivity (Fig. S10B), all units in each layer of TDANN and non-topographic control models were used to compute the predictivity score on the macaque IT neurons.

### Viewpoint invariance

To evaluate the emergence of viewpoint invariance across successively higher level TDANN face patches, we investigated the distributions of facial identity and viewpoint tuning in model layer cIT and aIT. For this analysis, we followed the same procedure described in (19): identity tuning half-widths and head orientation tuning depths were measured from responses of face-patch units to the face images with 25 identities and 8 head orientations (Fig. S8A). The width at half maximum of identity tuning was computed by sorting the responses to the 25 identities for each unit and identifying the smallest index among the ones whose responses are greater than half of the maximum response. Head orientation tuning depth was measured using face images at front view and left, right, up and down views at full profile. For each cell, the preferred orientation, giving the largest mean response, was identified. Then, the tuning depth was computed using the mean response to frontal faces (*R*_*frontal*_) and the mean response to full profile faces in the preferred orientation (*R*_*profile*_) as below:

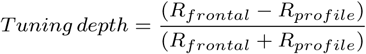

To compare with the statistics reported in (19), we computed mean value of identity tuning widths and standard deviation value of head orientation tuning depths using 20 randomly-initialized models (Fig. 6C). The distribution of tuning widths and tuning depths were plotted with a pool of face-patch units in all 20 models (Fig. S8B,C).

## ACKNOWLEDGMENTS

This research was supported by Samsung Scholarship (H.L.), Lore Harp McGovern Graduate Fellowship (H.L.), Sir Henry Wellcome Postdoctoral Fellowship (206521/Z/17/Z, K.M.J.), Intelligence Advanced Research Projects Agency (IARPA), US National Eye Institute grants R21-EY025863, the Simons Foundation (SCGB [325500, 542965], J.J.D), the Semi- conductor Research Corporation (SRC) and Center for Brains, Minds and Machines (CBMM, J.J.D.).

The authors declare no conflict of interest.

