## Supplementary Information for "Topographic deep artificial neural networks reproduce the hallmarks of the primate inferior temporal cortex face processing network"

**This PDF file includes:**

Supplementary text
Figs. S1 to S14
SI References

### Supporting Information Text

**Wiring cost.** To investigate whether the spatial correlation rule is a proxy for an axonal wiring cost minimization rule, we estimated the wiring cost in TDANN model networks and compared it with the non-spatial control ANN models. Conceptually, for a network like the ventral visual stream, axonal neural wiring includes the total length of feedforward axons (e.g. from pIT to cIT), the total length of feedback axons (e.g. cIT to pIT), and the total length of axons needed within each area (e.g. within pIT + within cIT). All of these costs will be explored in future work (Jozwik et al., unpublished).

In the present work, we focused on a simple model of the wiring length needed within area cIT, that we refer to as the total feedforward “branching” wiring length. The basic conceptual idea is this: when feedforward information is carried by neural spikes from a lower area (e.g. pIT) to a higher area (e.g. cIT), an axon carrying the spikes from each pIT "neuron" must at least travel the distance between the two areas and then branch to connect to each of the cIT "neurons" that it must make contact with. Note that the distance between pIT and cIT is the same in the baseline ANN models and the TDANNs. However, because the TDANN spatial correlation rule tends to spatially organize the cIT neurons according to response similarity, that suggests that the input branches (from each pIT) might not need to travel as far within cIT (relative to a random spatial layout). Indeed, this was the original intuition for the spatial correlation rule (we also note that recurrent connectivity within cIT might be reduced for the same reason, but the current ANN models are feedforward only, so they do not incur such wiring costs).

More specifically, to estimate the total branching wiring length within each model’s cIT, we performed the following procedure:

First, for each model pIT unit, we identified which cIT units it must connect to. Because the models start out as "fully connected" (all model pIT units (CONV5) connect to all model cIT units (FC6)), we pruned away all such connections that are not needed to maintain model function. To do this pruning, we ranked the pIT-to-cIT weights by absolute value. We then applied a threshold, setting all weights below that threshold to zero (effectively "pruning" the connection – no axonal connection needed). For each such pruned model, we determined its ImageNet validation performance. For each model network, we then determined the maximal amount of pruning (i.e. the maximal threshold) that could be done while dropping the performance by no more than 1% and by no more than 10%. The result was a pruned connectivity matrix (pIT-to-cIT) for each TDANN model network and for each baseline model network (cIT units were randomly arranged in the tissue map for the baseline model).

Second, we assumed that each pIT unit supplied one axon to the cIT and we determined the spatial location within cIT. We took this "arrival" location to be the location that pIT axonal arrival would tend to lead to the (approximate) lowest branching wiring length (within cIT) in its reaching to all the target cIT units that it must connect to (i.e. pIT-to-cIT connections that were not pruned for this pIT axon). Specifically, we defined the pIT axonal arrival location in cIT as the median location (x’,y’) of its target cIT units (x’ = median over target cIT x locations, y’= median over target cIT y locations).

Third, we computed the total axonal branching length needed for each pIT axon. We assumed one branch axon for each target cIT connection, which meant that the total axonal branching length was simply the sum of the Euclidean distances from its arrival location (above) to all of its cIT target neurons. We then computed the total (within cIT) axonal branching length for the overall model as the sum of the axonal branching lengths of each pIT unit.

After computing the total axonal branching wiring length in each model’s cIT (as defined above), we found that TDANN cITs have 6% and 19% lower branching wiring length compared to the non-spatial control models (numbers are for 1% and 10% ImageNet performance drop, respectively). In addition, when evaluating the ImageNet performance drop with pruning, we found that the same number of removed weights yielded a smaller performance drop in TDANN cITs than in the non-spatial control models, suggesting that TDANNs are less sensitive to the pruning of connections. In sum, our results thus far suggest that TDANN models derived via the spatial correlation rule (see Figure 1 and Materials and Methods) have modestly lower wiring costs than baseline models, even with nearly identical levels of performance.

**Reliability of face selectivity across different test stimulus sets.** To investigate how robust the face selectivity of TDANN model units is across different test sets, we computed the reliability of the spatial pattern of face selectivity across the test sets. More specifically, we first computed the face selectivity (d’) values over all TDANN cIT model units using each of the four test sets (1–4). Then, we measured the reliability score as Pearson correlation between the d’ values for each of six possible test set pairs. Figure S14A shows the reliability scores of the baseline TDANN model cITs, where each column corresponds to each of the test set pairs.

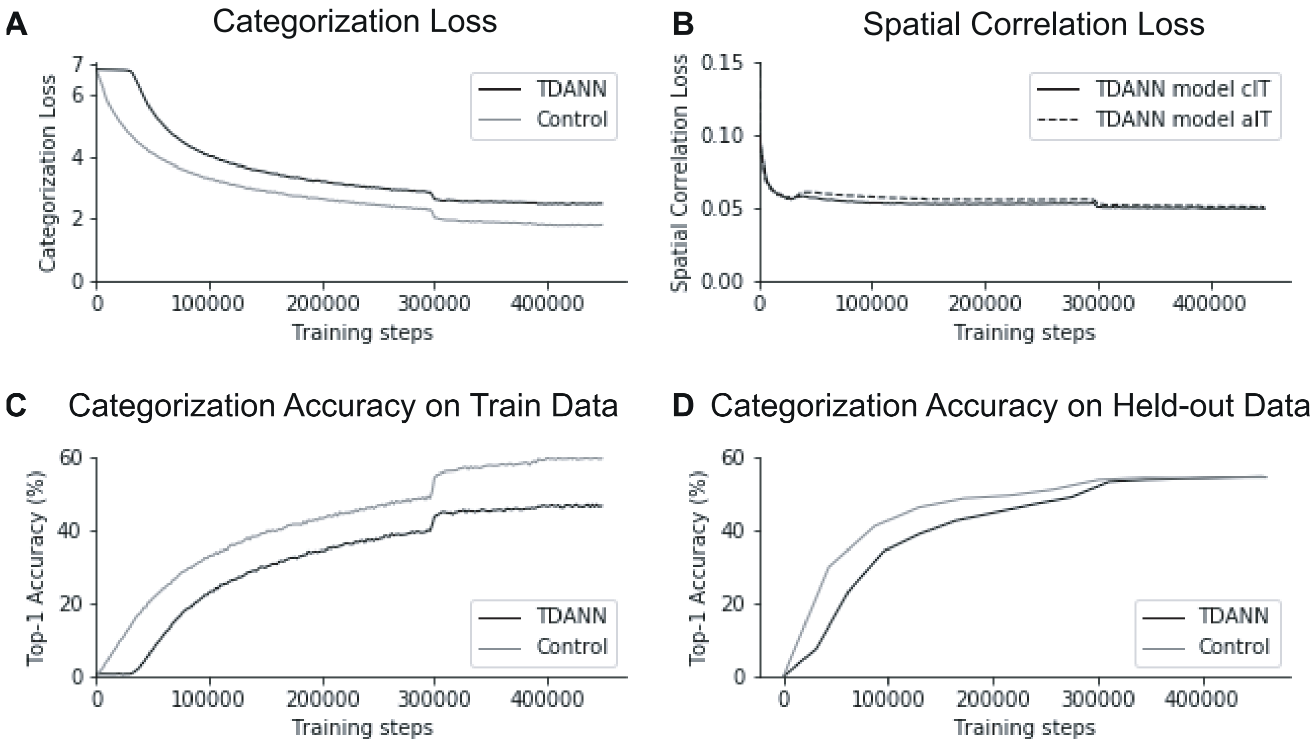


**Fig. S1.** Training results from example TDANN and non-topographic control models. A) Overall categorization loss of an example TDANN model (black) and an example non-topographic baseline control model during training (grey; same network architecture, but synaptic weights optimized for categorization loss only). The x axis represents the training step, where each step corresponds to a batch of 256 training images. B) Spatial correlation loss of the same TDANN for its model cIT (solid line) and model aIT (dashed line) throughout the training. C & D) Categorization performance (top-1 accuracy) of the TDANN model and the non-topographic control model on the training images (C) and held-out validation images (D). While there is a difference between the final categorization accuracy of the TDANN model and then non-topographic control model as measured on the training images, the TDANN model shows nearly the same validation accuracy as the non-topographic control model.

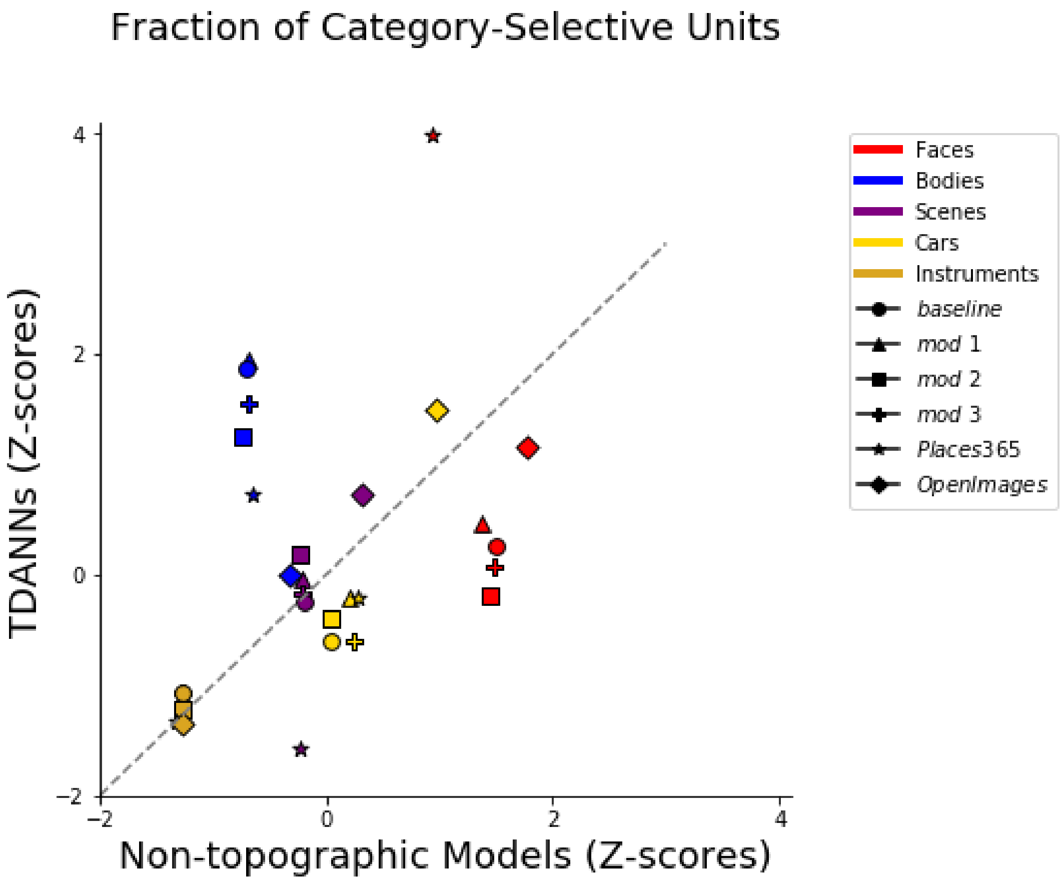


**Fig. S2.** Fraction of category-selective units ($d' > 0.85$) in TDANN model cITs vs non-topographic model cITs trained with different visual diets (see Figure 7 for details). The color and shape of the dots indicate the category of interest and the visual training diet, respectively. To neutralize any variability due to different image test sets, the fraction values were z-scored for each test set, and then averaged across three different test sets (1, 2, 5). Each dot indicates the value averaged over the randomly-initialized models first, then averaged over the three test sets.

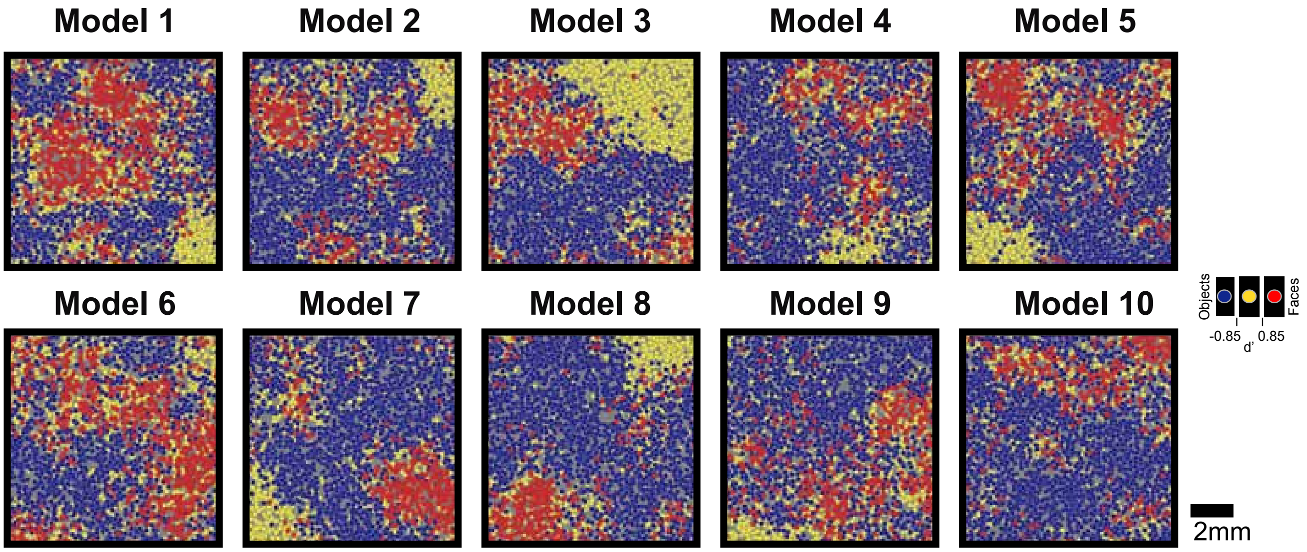


**Fig. S3.** Face selectivity maps of ten TDANN models cIT using the test set from (1). Face-selective units ($d' > 0.85$) are shown in red; non-face object-selective units ($d' < -0.85$) are shown in blue; units that are not strongly biased in either direction ($-0.85 \leq d' \leq 0.85$) are shown in yellow. Each map corresponds to each of the ten differently initialized models. Note the presence of one or two face patches in each model cIT, but precise tissue location varies within the 10 mm range. Scale bar: 2mm.

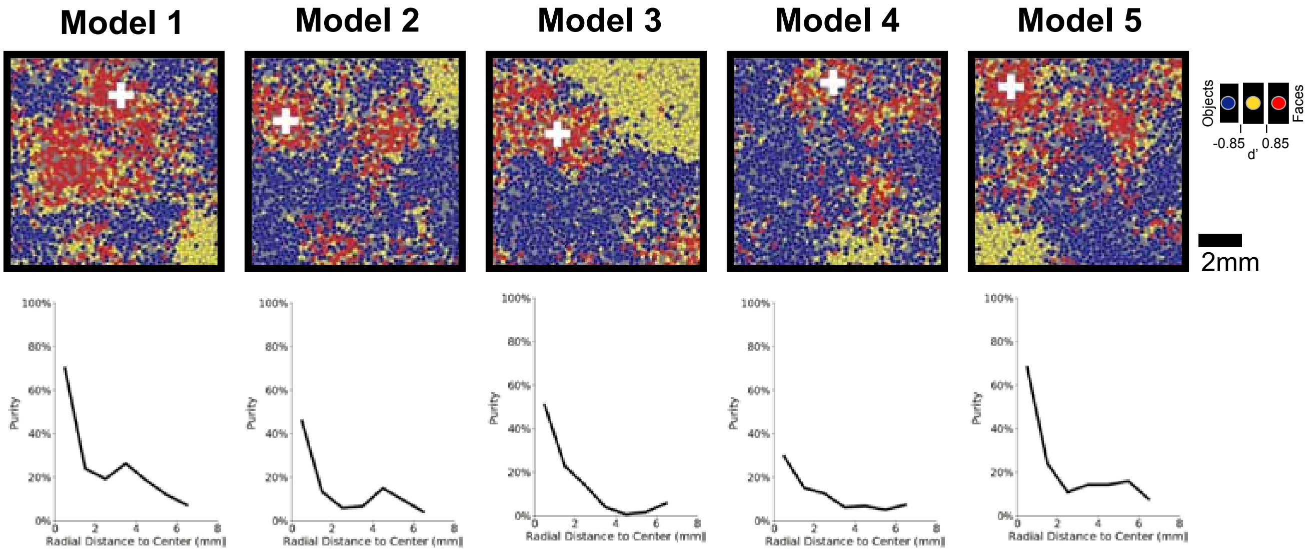


**Fig. S4.** Spatial profile of face patches in TDANN model cIT. Upper panels show the face selectivity maps that are shown in Figure S3 as well. Lower panels show the purity of face selectivity as a function of distance from the center (shown as a white plus marker in each map) of the dominant cIT face patch, corresponding to the maps immediately above. Each column corresponds to each of the differently initialized models, and Figure 3B (rightmost) shows the mean over all 20 models. Scale bar: 2mm.

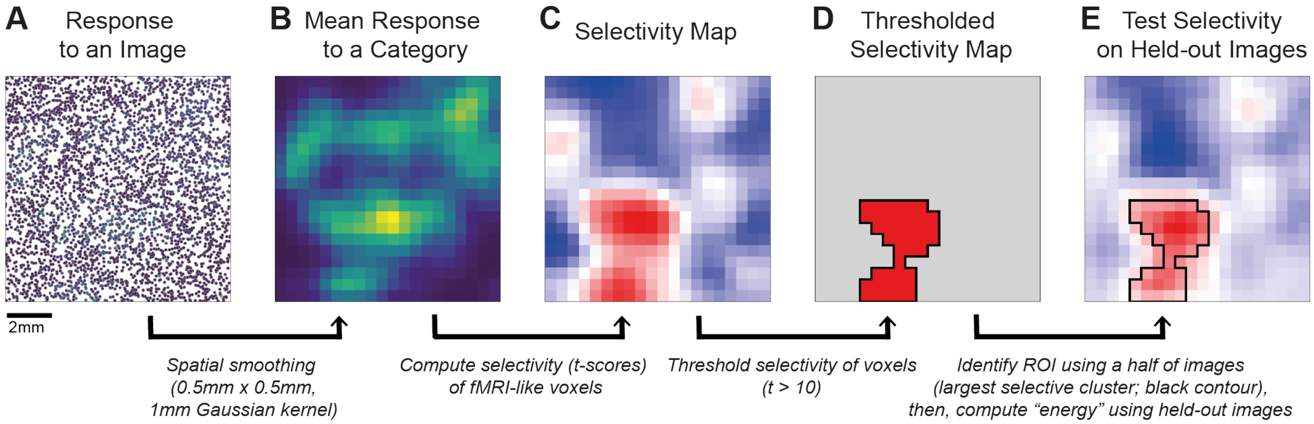


**Fig. S5.** Illustration of procedures to generate a simulated fMRI category selectivity map and measure the cluster strength. A) Raw activation of TDANN model cIT units to a single image. Each dot corresponds to each unit. B) Smoothed activation map in a per-voxel basis. To simulate the fMRI-like images, the voxel responses were computed for every 0.5mm x 0.5mm spatial bin ("voxel") by spatially smoothing the image-driven activation values of units with a 1mm Gaussian kernel (full width at half maximum). This is taken as the underlying voxel response map for that image. C) The category selectivity of each voxel is computed as $t$ scores from the voxel response maps (A) resulting from all images in the test category and all images in the anti-category (see Materials and Methods). D) Thresholded selectivity map is all voxels that have a selectivity value (t) greater than 10 (i.e. corresponding to $p < 0.001$). The region of interest (ROI) of the test category is taken as any group of connected voxels in that thresholded selectivity map (Black line). The primary ROI was taken as the spatially largest such region. E) The cross-validated (aka test) selectivity map (a contrast of two sets of images) is computed from held-out images of the test category and held-out images from the non-category. The "cluster strength" was computed as the sum of these cross-validated selectivity intensities of all voxels in the primary ROI. The example maps shown are for a single TDANN model cIT, and the test category was faces (with non-faces as the contrast image set) using the test set from (2). For cluster strength results (e.g. Figure 4 and 7), the identical procedure (above) was applied for each TDANN model, each test category, and each set of test images. Scale bar: 2mm.

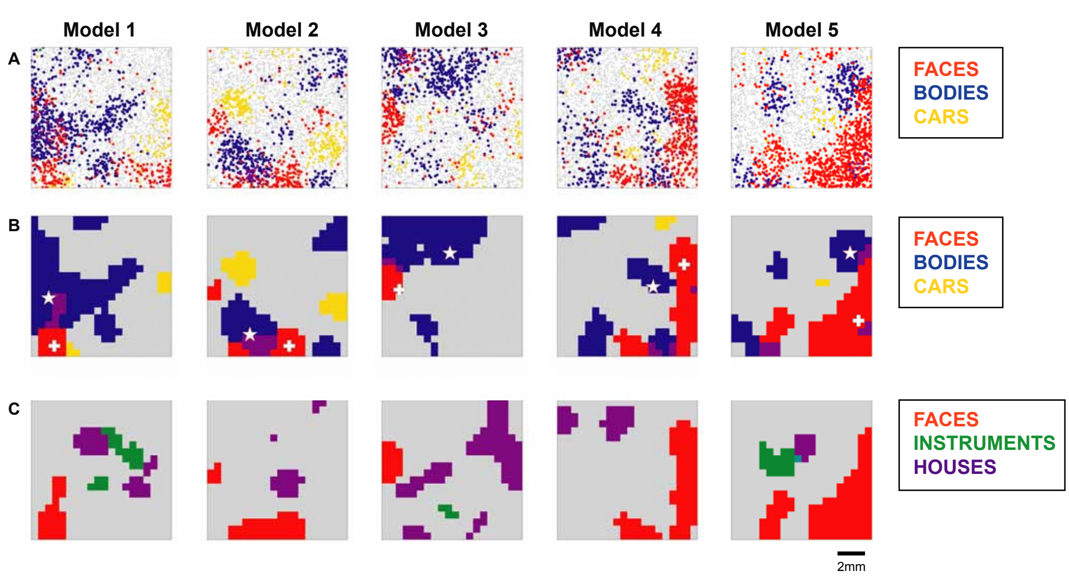


**Fig. S6.** Selectivity maps over different categories in five example TDANN model cITs, using the test set from (2). A) Each map shows superposition of selectivity ($d' > 0.85$) for faces (red), bodies (blue), and cars (yellow). (Analogous to 4A (upper) in the main text.) Units that were selective for both faces and bodies are shown in purple. B) Simulated fMRI selectivity maps (Analogous to Figure 4A (lower) and 4B in the main text.) Each plot is a simulated fMRI selectivity map for the indicated categories. The white-colored plus and star markers indicate the locations of peak selectivity for faces and bodies, respectively. C) Same as panel B, but with different categories: faces (red), houses (purple), and instruments (green). Each column corresponds to a differently initialized model. Scale bar: 2mm.

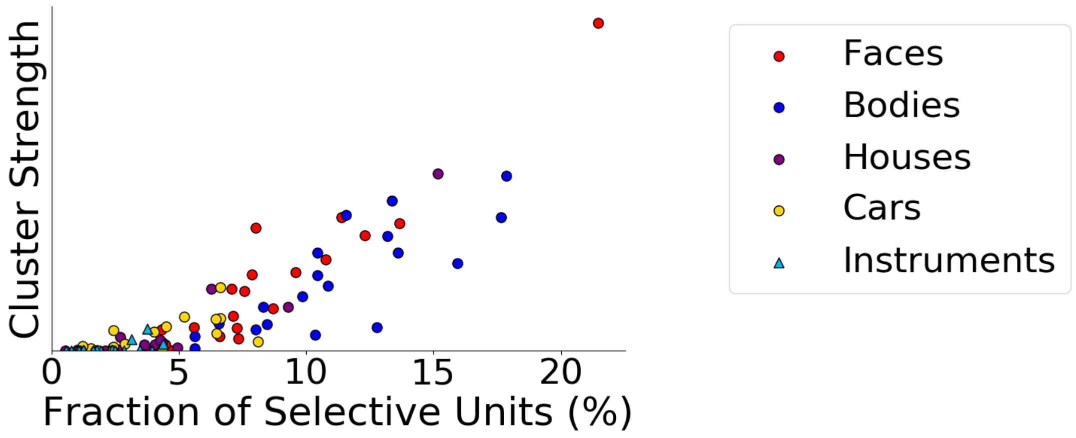


**Fig. S7.** Fraction of category-selective units ($d' > 0.85$) versus cluster strength of category-selective voxels ($t > 10$) in fMRI simulated selectivity maps (i.e. Figure 4A (lower) and 4B), in response to the test set from (2). Each dot indicates results from one of the 20 randomly-initialized TDANN models, and the color/shape of the dot indicates the tested category.

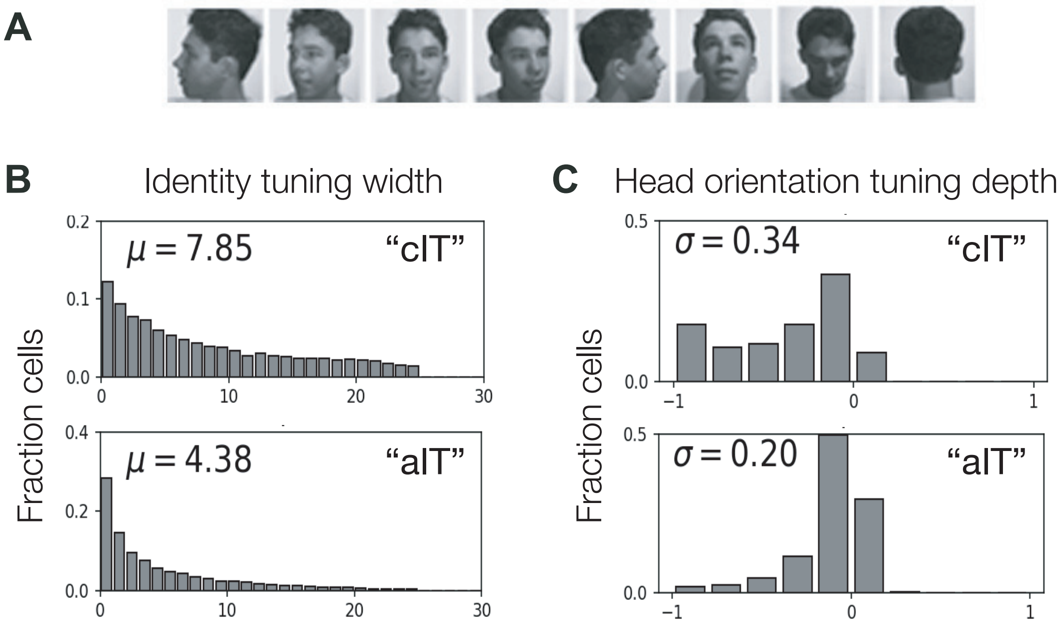


**Fig. S8.** Facial identity and viewpoint tuning of TDANN face patches in model cIT and model aIT, following the conventions of (6). A) One of the 25 face identities from the image set is shown at eight different viewpoints (adpated from (6)). B) Distribution of the identity tuning widths for neurons sampled from TDANN cIT (upper) and TDANN aIT (lower) face patches. Smaller tuning half-widths indicate greater identity selectivity. C) Distribution of head orientation tuning depths in TDANN face patches. Tuning depths close to 0 indicate greater viewpoint invariance. The mean of identity tuning widths ($\mu$) and standard deviation of head orientation tuning depths ($\sigma$) are shown in each plot and correspond to the bar heights in Figure 6C. The distributions were drawn from all face patches in 20 randomly initialized TDANN models. The distributions of empirically observed values for these measures for neurons in macaque face patches can be found in (6).

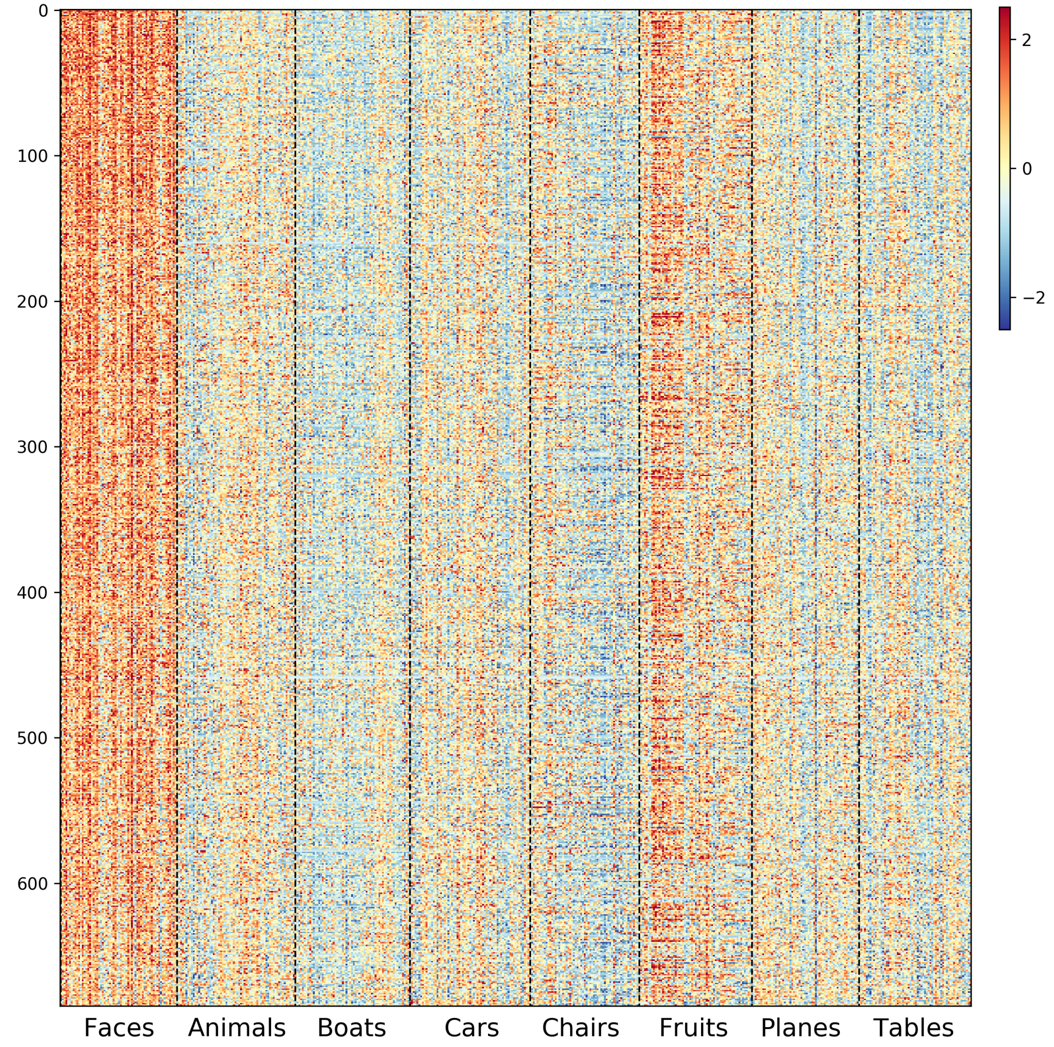


**Fig. S9.** Responses of TDANN model cIT face-patch face-selective units ($n = 685$). Color intensity indicates the model unit's responses to each of 501 images from eight different categories in (7). The responses of each model unit were standardized as z-scores over held-out test images.

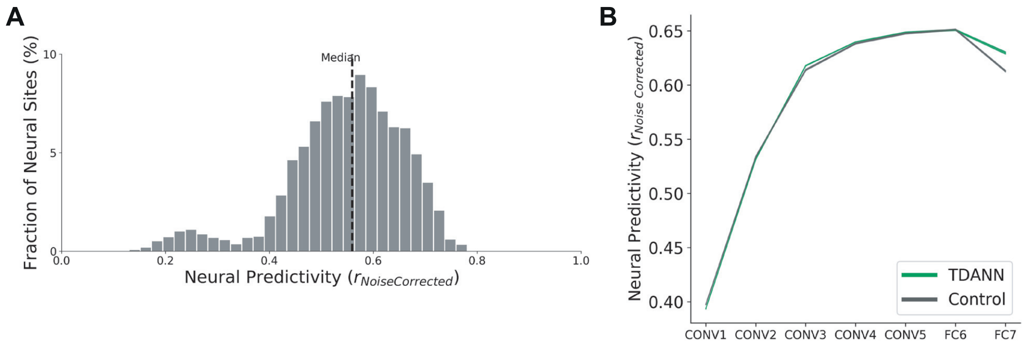


**Fig. S10.** Model predictivity of the image by image pattern of responses of macaque IT neurons. A) Distribution of TDANN model predictivity scores over 34 macaque IT face-selective neurons. Here the test images are the 501 images used in (7). The predictivity score for each IT neuron was computed as the noise-corrected Pearson correlation between IT neural responses and the TDANN model predicted response pattern of that neuron using held out images (here, the model predictions are based on the regression of face-selective model units in each model cIT face patch; see Materials and Methods for details). The distribution shows predictivity scores for all twenty TDANN models (with random initializations) over all 34 monkey IT face-selective neurons, and the dotted line corresponds to the median score. B) Median IT neural predictivity for each layer of TDANN (green) and non-topographic control (grey) models. Here, the median fit is computed across all 168 IT neurons, then averaged over 20 (TDANN) or 10 (non-topographic control) randomly initialized models to yield the plotted lines. The shaded area around each dark line (barely visible as it is less than 0.01) represents the SEM across all randomly initialized models.

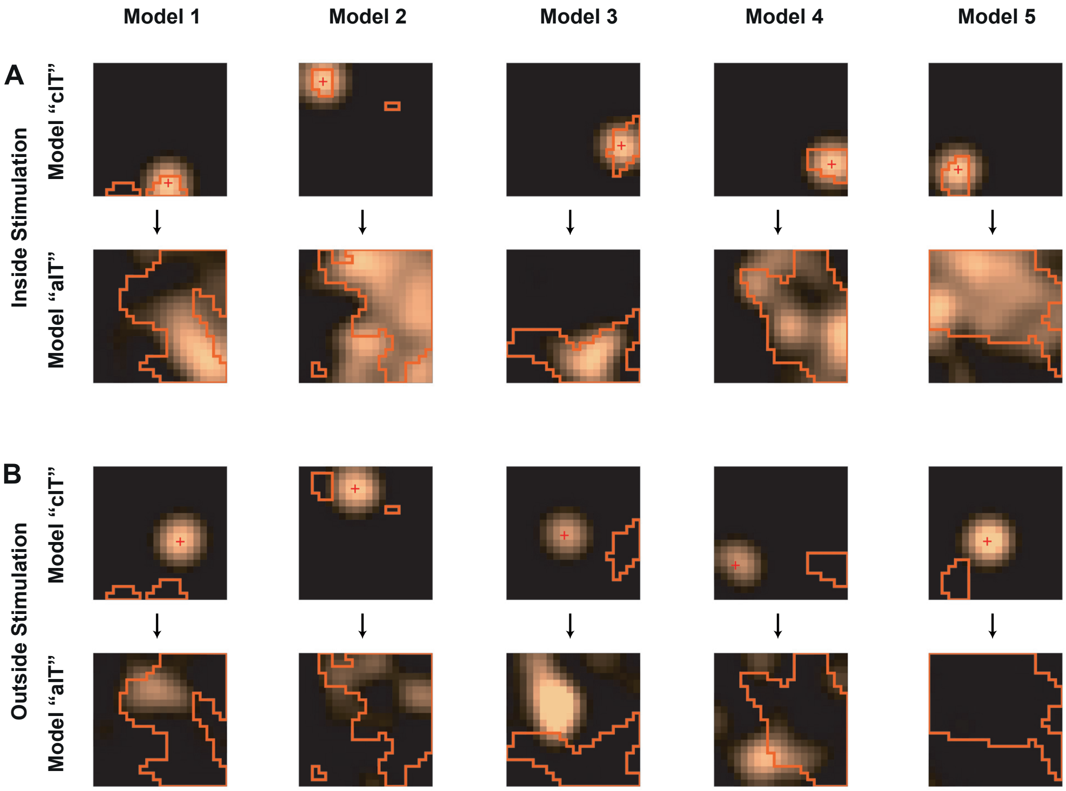


**Fig. S11.** Additional predicted electrical stimulation results simulated in model cIT of five example TDANN models. Orange lines demarcate the boundaries of face patches (see Figure 5 for procedure to derive). (A) shows cIT stimulation inside a model cIT face patch, and (B) shows stimulation outside cIT face patches. Each pair of panels shows the simulated model cIT stimulation (upper) and the observed responses in model aIT (lower). The red plus marker in the model cIT map indicates the simulated stimulation site and the color map (black to bright orange) represents the activation levels of the voxels in each layer.

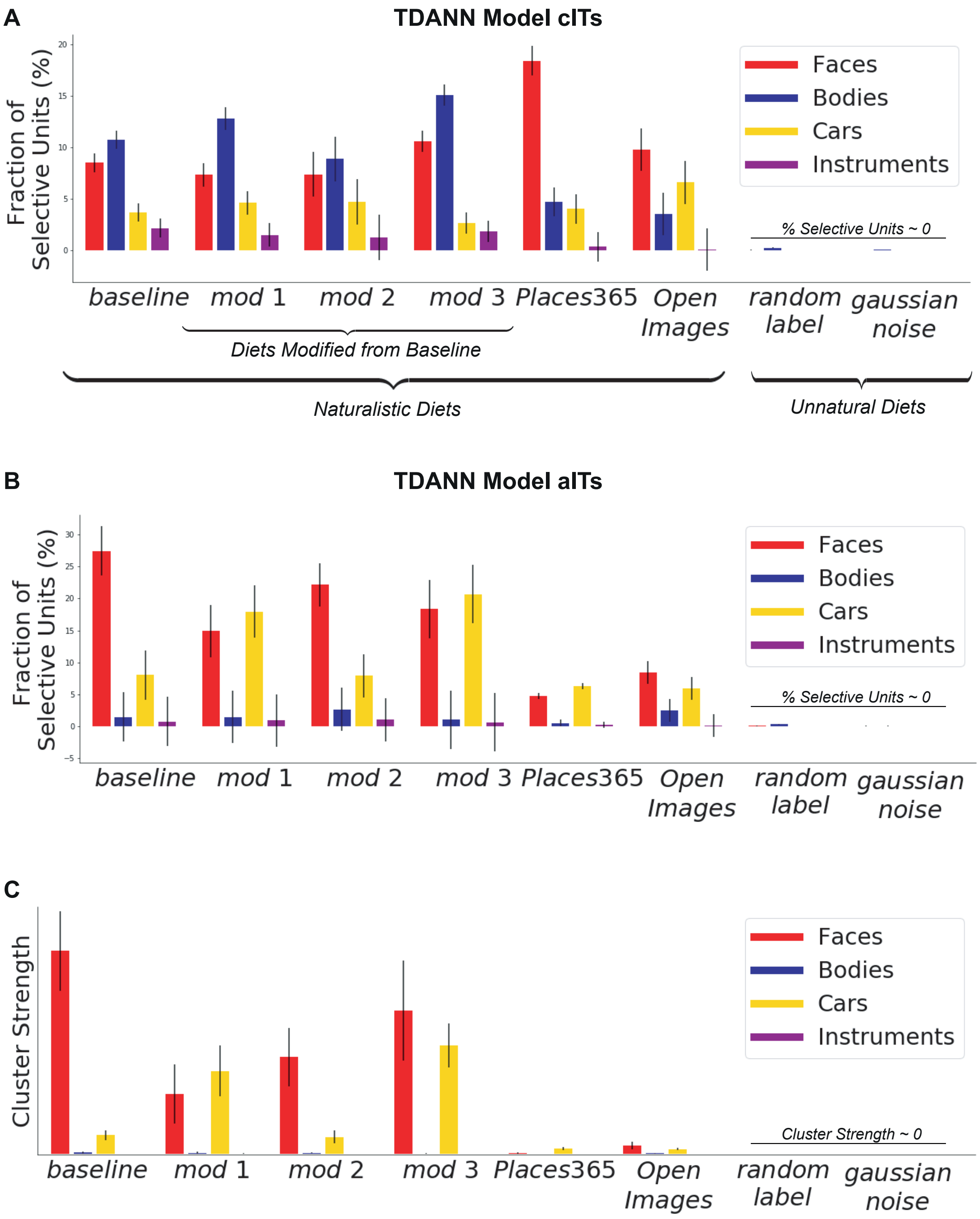


**Fig. S12.** Additional summary statistics of the effect of visual diet on category selectivity in TDANN model cITs (upper) and model aITs (lower two). Same as Figure 7, the different training diets are: the baseline diet, three additional diets with different categorical distributions modified from the baseline (*modification 1-3*; Figure 7A), Places365 (8), Open Images (9), ImageNet (10) images with randomly shuffled labels (*random label*), and random Gaussian noise images (*gaussian noise*), from left to right. The first two plots show the fraction of category-selective units ($d' > 0.85$) in TDANN model cITs (A) and model aITs (B). Panel C shows cluster strength of category-selective voxels ($t > 10$) in TDANN model aITs (as defined in Figure 4). Red indicates selectivity for faces; blue for bodies; yellow for cars; purple for instruments. Each bar height indicates the mean value and error bars indicate the SEM over 20 (for baseline diet) or 10 randomly-initialized models.

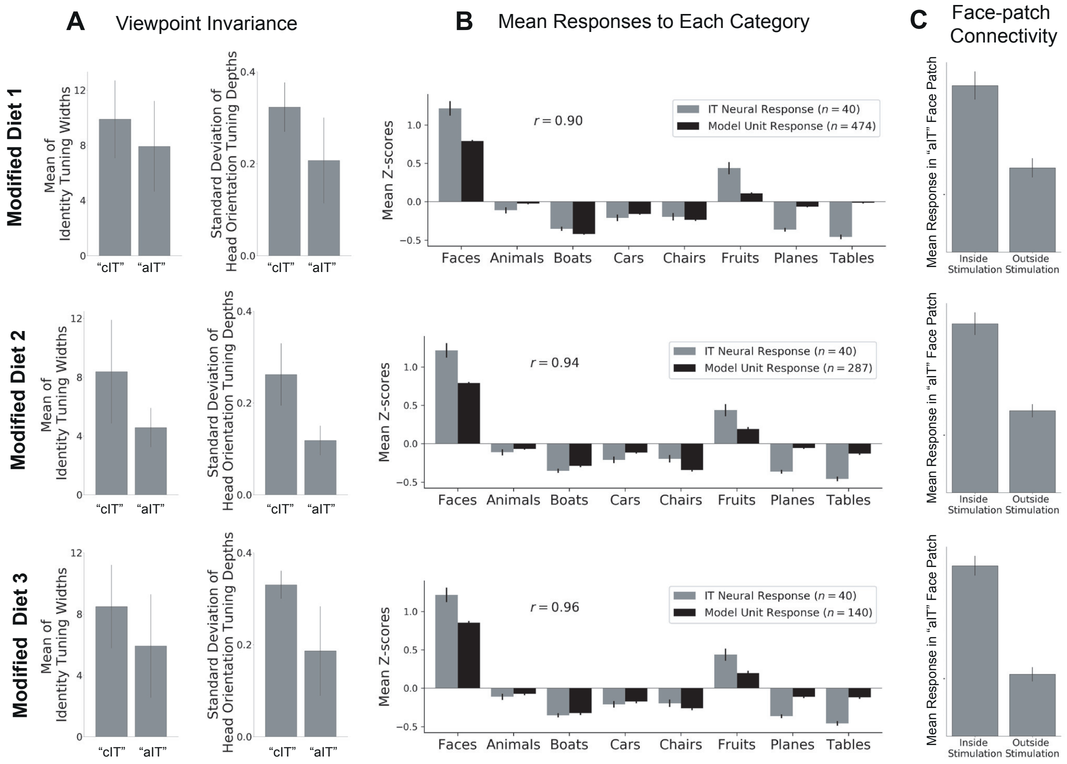


**Fig. S13.** Additional hallmarks of IT face processing network were robustly found regardless of the visual experience (training) diets. Each row corresponds to each training visual diet in Figure 7A. A) Facial identity (left; mean identity tuning widths) and viewpoint tuning (right; standard deviations of head orientation tuning depths) in TDANN face patches. Each bar represents model cIT and model aIT, respectively. Computed as in Figure 6C of main paper. B) Average response to different categories across macaque face-selective neurons (gray, $n = 61$, (7)) and face-selective units from model cIT face patches (black, $n = 685$). Computed as in Figure 6B of main paper. C) Activation of the model aIT face patch units for stimulation inside (left) and outside (right) of model cIT face patches. Each bar height indicates the mean value and error bars indicate the SEM over 20 randomly-initialized models. Computed as in Figure 5 of main paper.

**A**

**B**

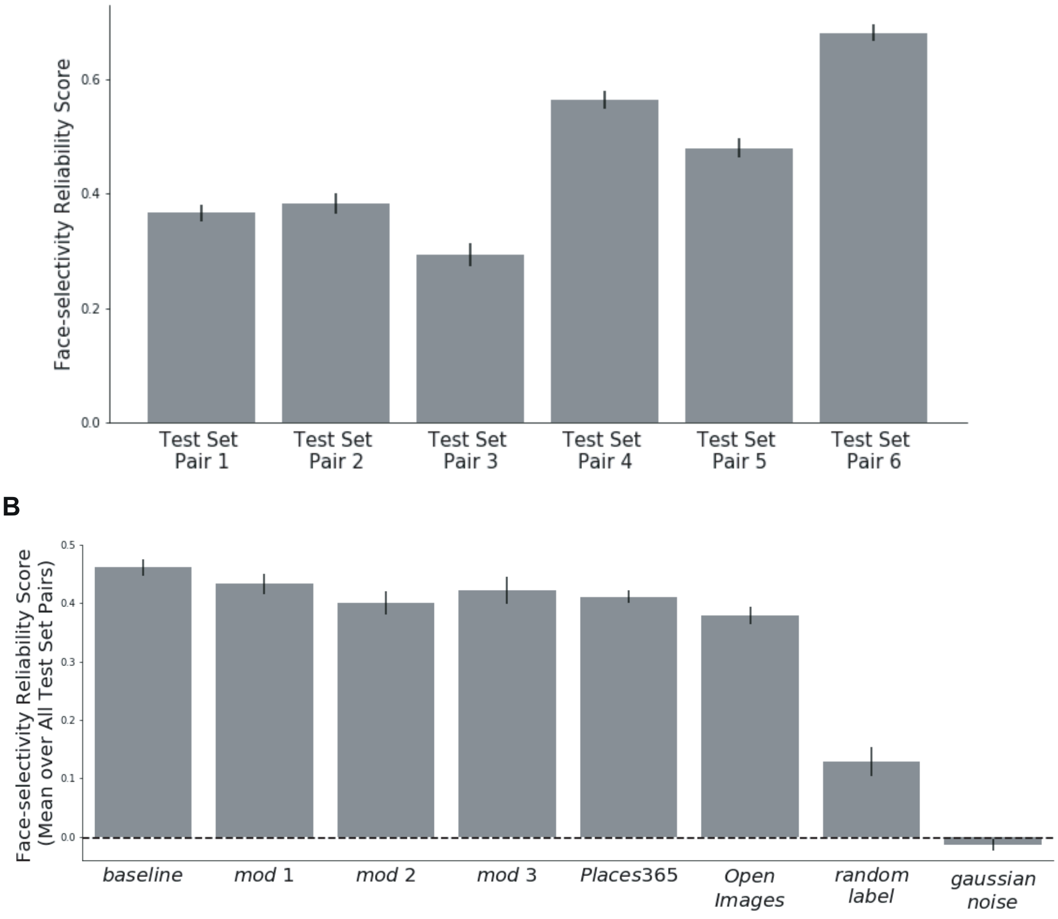


**Fig. S14.** Reliability of the spatial pattern of face selectivity in TDANN models. A) Face-selectivity reliability scores of baseline TDANN models computed over different pairs of image test sets: (1, 3), (1, 2), (1, 5), (2, 3), (3, 5), (2, 5), from left to right. Each reliability score was computed as the Pearson correlation of $d'$ values (of all model cIT units) as assessed by each pair of image test sets. Each bar height indicates mean value and error bars indicate the SEM over 20 randomly-initialized models. B) Mean face-selectivity reliability scores of TDANN models trained with different training diets: the baseline diet, three additional diets with different categorical distributions modified from the baseline (*modification 1-3*; Figure 7A), Places365 (8), Open Images (9), ImageNet (10) images with randomly shuffled labels (*random label*), and random Gaussian noise images (*gaussian noise*), from left to right (see Materials and Methods for details). Each reliability score was averaged over all possible pairs of test sets used in panel A, then averaged over differently initialized models. Each bar height indicates the mean value and error bars indicate the SEM over 20 (for baseline diet) or 10 randomly-initialized models.
